# Open Benchmarking for Cell-Based Multiscale Models: Lessons from a Community Initiative

**DOI:** 10.1101/2025.07.16.664358

**Authors:** Thaleia Ntiniakou, James Osborne, Jieling Zhao, Jules Dichamp, Nicolò Cogno, Randy Heiland, Kwabena Amponsah, Jiri Pesek, Tobias Duswald, Othmane Hayoun-Mya, Jack Jennings, Matteo Pedrazzi, Ryan Bournes, Marco Ruscone, Fergus Cooper, Vincent Noël, Matthew I Leach, Alejandro Madrid-Valiente, Jose Estragués-Muñoz, Adam Smelko, Taghi Aliyev, Marco Manca, Joe Pitt-Francis, Gary Mirams, Paul Macklin, Alexander G. Fletcher, Vasileios Vavourakis, Stefan Hoehme, Jose Carbonell-Caballero, Roman Bauer, Van Paul Liedekerke, Dirk Drasdo, Alfonso Valencia, Arnau Montagud

## Abstract

The emergence of virtual human twins (VHT) in biomedical research has sparked interest in multiscale *in silico* modelling frameworks, particularly in their application bridging cellular to tissue levels. Among the diverse array of multiscale modelling tools, off-lattice center-based agent-based models (CBM) offer a promising approach due to their depiction of cells in 3D space, closely resembling biological reality. Despite the proliferation of CBM tools addressing various biomedical challenges, a comprehensive and systematic comparison among them has been elusive.

This paper presents a community-driven benchmark initiative aimed at evaluating and comparing CBM for biomedical applications, akin to successful efforts in other scientific domains such as the Critical Assessment of Protein Structure Prediction (CASP). Enlisting developers from leading tools like BioDynaMo, Chaste, PhysiCell,TiSim, and CompuTiX, we devised a benchmark scope, defined metrics, and established reference datasets to ensure a meaningful and equitable evaluation. Unit tests targeting different solvers within these tools were designed, ranging from diffusion and mechanics to cell cycle simulations and growth scenarios. Results from these tests demonstrate varying tool implementations in handling diffusion, mechanics, and cell cycle equations, emphasising the need for standardised benchmarks and interoperability.

Discussions among the community underscore the necessity for defining gold standards, fostering interoperability, and drawing lessons from analogous benchmarking experiences. The outcomes, disseminated through a public platform in collaboration with OpenEBench, aim to catalyse advancements in computational biology, offering a comprehensive resource for tool evaluation and guiding future developments in cell-level simulations. This initiative endeavours to strengthen and expand the computational biology simulation community through continued dissemination and performance-oriented benchmarking efforts to enable the use of VHT in biomedicine.

## Introduction

Virtual Human Twins (VHTs), a “virtual representation that serves as the real-time digital counterpart” of a patient, have sparked interest in the last few years for their potential to deliver a truly personalised treatment to patients, as they are “designed to accurately predict specific quantities that, while difficult or impossible to measure directly, would be very useful to support a specific clinical decision” (Viceconti et al., 2024). The potential of VHTs is rooted in recent advances in numerical and theoretical methodologies as well as computing infrastructures that enable more comprehensive and tailored approaches to a vast array of biomedical topics (Cogno et al., 2024; Montagud et al., 2021). These applications include drug repurposing (Ponce-de-Leon et al., 2023), mechanistic understanding of complex interdependent processes in disease progression (de Montigny et al., 2021; Hoehme et al., 2023) including treatment response to cancer (Demetriades et al., 2022; Gazeli et al., 2022), and the optimisation of drug delivery (Ponce-de-Leon et al., 2022) to maximize the therapeutic effects of multimodal treatments (chemo, radio, immunological) (Cogno et al., 2024; Hadjicharalambous et al., 2022; Ioannou et al., 2024; Mongeon et al., 2024; Wang et al., 2024), supporting patients throughout their clinical pathway.

Agent-based modelling stands apart from other modelling methods by simulating complex systems based on the interactions among individual elements constituting the system’s population and, in doing so, by bridging diverse temporal and spatial scales. In the context of biology, agent-based models (ABMs) can simulate proteins, bacteria, biological cells, individual organisms, or any identifiable entities. ABMs can describe the collective behaviour arising from these entities’ actions and produce system-level outcomes. This computational methodology has demonstrated predictive capability in modelling biological systems at the cellular level, successfully replicating complex interaction networks across varying spatiotemporal scales (An et al., 2009; Cogno et al., 2024; de Montigny et al., 2021; Van Liedekerke et al., 2015). Such systems exhibit nonlinear behaviour, stochasticity, and varying degrees of interconnection between entities (Metzcar et al., 2019). While most individual components of an ABM are at least partially amenable to analytical solutions, the combined system typically is not.

Their ability to bridge scales makes ABMs an excellent complement to *in vitro* and *in vivo* experimental methods, which often have limited resolution capacity across biological scales and can be costly in terms of time and resources. Indeed, ABMs are widely used as multiscale modelling tools for biological tissues, as they can integrate various methods to simulate processes occurring at different time scales - such as diffusion, mechanical interactions, and cell signalling - as well as at different spatial scales, ranging from protein functions to tissue and organ dynamics (de Montigny et al., 2021; Schaffer & Ideker, 2021). These capacities have allowed ABMs to be considered as potential candidates to build VHTs and a means towards reaching the principles of the 3Rs (Replacement, Reduction and Refinement) of animal research (Dichamp et al., 2023; Jean-Quartier et al., 2018). In fact, simulations are rapidly gaining prominence and replacing animal models in the initial phases of drug development, ranging from screening campaigns to identify novel drug targets and lead compounds to safety and efficacy assessments (Wadman, 2023).

ABMs can vary widely in their algorithmic and numerical formulation. For an exhaustive overview of agent-based modelling methods, refer to review articles (Metzcar et al., 2019; Osborne et al., 2017; Van Liedekerke et al., 2015). Several approaches commonly used in computational biology include the following:

1. **Center-Based Models (CBM):** CBM represent cells as simple geometric objects (usually spheres) whose centers move in response to forces in continuous 3D space without grid constraints. These models allow us to understand complex systems dynamics but ignore the detail of every cell’s shape (Osborne et al., 2017).
2. **Voronoi or Boundary-based models:** These are a type of boundary-tracking models that use Voronoi tessellation to define the complex boundary of center-based objects (such as those in a CBM) and can be used to understand the physical constraints that drive the pattern-like organisation of biological tissues, especially in confluent structures (González-Valverde & García-Aznar, 2017).
3. **Vertex Models (VM) and Deformable Cell Models (DCM):** Both model types allow to simulate complex cell shape by tracking the boundaries of cells using multiple vertices per cell. This permits them, for instance, to compute subcellular forces enabling to address mechanobiological questions (Fletcher et al., 2014; Van Liedekerke et al., 2019). In comparison to CBM, they allow simulations in more detail but are more computationally expensive to execute.
4. **Cellular Potts models (CPM):** These are lattice-based models in which cells are represented as a set of small domains that can move on a fixed grid. The dynamics is controlled by energy functions (Hamiltonians) governing adhesion, volume constraints, and interactions. As with VM and DCM, complex cell shapes can be modelled with this type (Merks & Glazier, 2005).

Each of these modelling methodologies has its unique strengths and applications, making them valuable resources in the field of multiscale modelling. They help understanding complex systems by providing a framework to integrate information across multiple spatio-temporal scales. Often, one needs to make a trade-off between higher detail (VM, DCM, CPM) and computational feasibility (CBM). Here we focus on CBMs as physically well-founded tools that have the ability to implement physical and biological functions as agents in a realistic continuous space in a straightforward and computationally inexpensive way. CBMs can represent biological cells as particles that can grow, move, differentiate, multiply and die in three-dimensional space while also interacting with other cells and their microenvironment; thus, CBM simulations can facilitate virtual representation of cells that closely mirrors biological reality.

In summary, CBMs 1) support representations of fully populated (i.e., cells in the epidermis) as well as sparsely populated spaces (i.e., cells in connective or stromal tissue), 2) allow the integration of spatio-temporal interactions (biomechanical and functional) between various tissue elements, 3) capture cell biomechanics with an acceptable realism (Drasdo & Hoehme, 2005, 2012), and 4) are computationally efficient tools when considering simulations of tissues of cubic centimeters with micrometer resolution [REF BioFVM-B preprint]. Other approaches can have some of these characteristics but none have them all.

Nowadays, many scientifically developed tools that use CBMs in biology are available in the community and can be deployed to study a broad spectrum of biomedical problems. Despite this, these tools were never thoroughly and systematically compared in terms of available functionalities and expected behaviours. Benchmark can be defined as the comparison of a set of tools running a set of tests used as standard points of reference that facilitate evaluating the performance and characteristics of these tools (Peters et al., 2018).

Although benchmarking is common throughout fields like bioinformatics, computational mechanics, and computational fluid mechanics, ABMs have been scarcely benchmarked (Koning & Gropp, 2024; Moreno et al., 2019). Importantly, the comparisons of specifically CBMs have been find the best tool or configuration for a specific task (Pleyer & Fleck, 2023) or comparing a newly developed tool against a few similar tools from the field. Notably, the need for VHTs is further motivation to fill this gap and present CBM tools that deliver simulations that have the same common functions signatures, or at least that the deviation of the results is methodologically-grounded.

To this end and as part of the PerMedCoE project [REF PerMedCoE consortium preprint], we have run a community-driven benchmark following successful benchmarking efforts, such as the Critical Assessment of Protein Structures Prediction (CASP) (Kryshtafovych et al., 2023), running since 1994; and other popular and widely-accepted benchmark competition in image analysis and AI, such as image classification benchmarks based on given datasets (Bassi et al., 2025; Zheng et al., 2024), or the regular medical segmentation benchmarks organized by the MICCAI conference (Bilic et al., 2023).

We define “community” as a diverse group of scientifically oriented people, working within the same field, in our case, the off-lattice cell-level simulation field inside the wider computational biology field. Despite varied backgrounds and experiences, these individuals face similar challenges and are united by a common goal: to collaborate and discover optimal solutions to these shared problems.

Interestingly, these benchmarking efforts serve multiple purposes and have different benefits to various stakeholders within the community (Nature Methods Editorial Board, 2014). For tool developers, these benchmarks provide an objective platform for comparing their tools against others in the field. This encourages the implementation of more efficient methods and the development of new tools, by highlighting challenging areas, blind spots, and untapped opportunities in existing methodologies.

For end-users, access to benchmark results is invaluable when selecting a tool for a specific task. This not only ensures that users are equipped with the most effective tool for their needs but also steers the user base toward the latest developments in the field. This, in turn, benefits the authors of the original tools, as it brings attention and users to their implementations.

Through this community-driven benchmark, we aim to foster a collaborative environment that promotes innovation and advancement in the field of multiscale modelling. We believe that this initiative will lead to more comprehensive benchmark practices and more efficient and effective tools, ultimately benefiting the broader scientific community.

Disclaimer: this document is the first version of a work in process and the benchmark tests here represent a minimal, foundational comparative framework, not a comprehensive evaluation of each tool’s full capabilities.

## Methods

### Off-lattice multiscale modelling tools that participated in the benchmark

The following is a list of tools that were benchmarked. All of them are off-lattice, centre-based, agent-based modelling tools that can simulate cells as spherical shapes. Refer to Table S1 for a comparative table of the tools and Supplementary File 1 for more details on the equations used by the models.

BioDynaMo (Breitwieser et al., 2021) (https://www.biodynamo.org/) is an open-source simulation tool that is fully parallelised, is able to offload computations to hardware accelerators while also load-balance agents and their simulated microenvironment (Breitwieser et al., 2023; Hesam et al., 2021). Being general-purpose, it allows simulating models from various fields by being extensible and modular, showcasing its use in neurite growth (Duswald et al., 2024), in cancer therapy (Demetriades et al., 2022; Gazeli et al., 2022) and in epidemiology examples. The code for BioDynaMo is available under an Apache-2.0 licence at: https://github.com/BioDynaMo/biodynamo. We used version 1.04 for developing the different tests.

Chaste (Cooper et al., 2020) (https://chaste.github.io/) is an open-source, general-purpose simulation package for modelling soft tissues and discrete cell populations, where select simulation paradigms can be parallelised with the MPI library. This tool allows using different agent-based modelling frameworks (centre-based or not) on a given problem, enabling users to select the most appropriate one for their research and to better understand the limitations of each one of them. Chaste has been used for a wide range of projects, such as intestinal (Dunn et al., 2013) or colonic crypt (Dunn et al., 2012) studies. The code for Chaste is available under a BSD 3-Clause licence at: https://github.com/Chaste/Chaste.

PhysiCell (Ghaffarizadeh et al., 2018) (https://physicell.org/), an open-source, flexible, off-lattice, agent-based framework for multiscale simulation of multicellular systems that currently support shared-memory parallelisation. The main advantage of PhysiCell is its lightweight, very efficient and self-contained framework. Additionally, PhysiCell can be expanded using add-ons, such as PhysiBoSS (Ponce-de-Leon et al., 2023), allowing the integration of individual Boolean models for the signalling networks embedded into each agent. The code for PhysiCell is available under a BSD 3-Clause at: https://github.com/MathCancer/PhysiCell. For developing the different tests, we used version 1.14.2 of PhysiCell.

TiSim (Hoehme & Drasdo, 2010a) (https://multicellular_modelling.gitlabpages.inria.fr/group_webpage/tisim/) is a source-available software that has been used for multicellular biological models, such as liver regeneration (Hoehme et al., 2010, 2023), introducing the deformability in CBM (Van Liedekerke et al., 2019) and the inclusion of signalling pathways controlling the agents’ behaviours (Ramis-Conde & Drasdo, 2012).

CompuTiX is a flexible open-source library for off-lattice agent based modeling (not only) in biomedical applications including liver tissue microarchitecture reconstruction with deformable cell model (Pesek J. et al, in preparation) and NMR signal modeling (Boulitrop C. et al., in preparation). The main focus of the library is on extensibility, flexibility and transparency of internal state. The source code is available under AGPLv3 only licence at https://gitlab.inria.fr/computix/computix.

### Infrastructure used to coordinate and disseminate the benchmark

We created a GitHub repository (https://github.com/PerMedCoE/observatory_benchmark) to host all running codes (when available), reference datasets, results files and scripts used to build the figures in this paper.

The different CBMs have been described in the PerMedCoE observatory of tools, which is a catalogue of tools part of the ELIXIR *bio.tools* registry, (https://permedcoe.bio.tools) and the benchmarks are publicly accessible at OpenEBench (Capella-Gutierrez et al., 2017) (https://openebench.bsc.es/projects/OEBC009) and at Zenodo (https://doi.org/10.5281/zenodo.15911390).

## Results

### Establishing a community-driven benchmark

As part of the PerMedCoE project observatory of tools, regular open calls were made to teams that had developed CBM tools with the goal of creating a benchmark that accurately reflects the challenges faced by the scientific community in terms of size, complexity, and content. After these calls, we gathered responses from the interested teams of BioDynaMo (https://www.biodynamo.org/), Chaste (https://chaste.github.io/), PhysiCell (https://physicell.org/) and TiSim (https://multicellular_modelling.gitlabpages.inria.fr/group_webpage/tisim/).

Developers from these tools assembled, and identified three key features to ensure a meaningful and fair benchmarking framework: the scope, the metrics and the reference dataset. The scope of the benchmark refers to putting down specific scientific questions that the tools should be able to address through simulations, i.e., defining *what* is going to be compared. The questions a benchmark can address are two-fold: technical, i.e., focusing on the tool’s performance (runtime, memory use, etc.), or scientific, i.e., focusing on the quality and accuracy of the simulation results within clearly defined tests. In the present benchmark, the first hackathon showed us that the tools presented differences in the mathematical models used and in the way they were implemented. Thus, we decided that the scope needed to be scientific, focusing on comparing how the tools’ simulate different very simple behaviours, and leaving the performance analysis for a future benchmark.

Further, the developers needed to define “*how”* to compare the tools. The selection of these metrics is critical to have a fair comparison and needs to be carefully chosen and by consensus among our community members. These metrics can be quantitative, like measuring the distance between each tool’s output and the reference result, or qualitative, when describing differences between tools’ methods. In the present benchmark, we compared the tools’ results among each other (either their dynamics or their steady states) and against analytical solutions or experimental data, whenever possible.

Finally, reference datasets are used as ground truth evidence or baseline data for the quantitative benchmark. These datasets need to be well defined, well documented, unbiased, and their use has to be supported by the community. They can be experimental data, analytical solutions or desired outcomes. In the present benchmark, we used experimental data from Brú and colleagues (Brú et al., 1998), we produced numerical solutions for the diffusion tests and a logistic growth dynamic model for the cell volume growth. These reference datasets are available at our GitHub repository: https://github.com/PerMedCoE/observatory_benchmark/tree/main/experimental_data.

### Defining and running tests to evaluate the core functions of the multiscale simulation tools

To benchmark all CBM tools, a set of tests were designed to study the ability for each of these tools to ensure reproducibility, to inspect how this was implemented and to interrogate the accuracy of these functions. These tests have been carefully selected for being simple, relevant and common to all tools: they can be used as building blocks of bigger simulations in cell biology, while also probing key modelling functionalities and can be implemented in all the tools. We refer to these tests as “unit tests”, when they probed one functionality of the CBMs, and “use case” when they combined the functionalities of several unit tests.

The unit tests list the simulation of (*a*) the diffusion of substances (e.g., cytokines, enzymes) that is the solver that is most frequently fired, with time steps as small as 0.01 minutes, (*b*) the biomechanics of cells that usually has time steps of 0.1 minutes and (*c*) the cell cycle evolution and growth, that usually has time steps of 6 minutes. Additionally, we designed a use case to replicate the growth experiment of an *in vitro* monolayer of a cell line. This use case combines cell mechanics and cell cycle growth functions.

The tools had so far been developed independently and had made diverging design choices about the inputs and setting needed to run simulations, thus a significant part of the initial efforts went into deciding how the same set of information translated for every simulation suite to launch with analogous starting conditions, as to limit bias in results due to involuntary tweaking or misalignment of initial conditions. Members of the teams gathered on September 22 and 23, 2022, at the Barcelona Supercomputing Center (BSC) in Barcelona, Spain, for a hybrid hackathon and benchmark the tools by running the defined unit tests. In the following months, we worked to ensure the unicity and completeness of the unit tests across all tools: that, in each test, all the teams implemented the same parameters and models when possible.

### Unit tests for the reaction-diffusion equation

The purpose of the diffusion unit tests is to quantify how different CBM codes perform in solving the diffusion equation (second law of Fick), and the related reaction-diffusion equation which includes source and sink terms.

The PDE solvers used by the codes allow to simulate concentration profiles of nutrients, growth factors and other chemical compounds in the extracellular space that play an important role in transient phenomena during stimulation of cell growth, differentiation, survival, inflammation, and tissue repair. In each simulation, the diffusion solver is called frequently and as such a proper estimation of its accuracy is essential.

All CBM codes tested include an implementation for solving the reaction-diffusion equation to model the balance of biochemical substances that diffuse, decay, or are secreted or uptaken by cells. Different numerical methods can be employed to solve for this equation, namely, the Finite Element Method (FEM), used by Chaste and TiSim, Finite Difference Method (FDM), used by BioDynaMo, and Finite Volume Method (FVM), used by CompuTiX and PhysiCell. Explicit-time Euler numerical scheme was used by BioDynaMo and Chaste and implicit-time Euler scheme by CompuTiX and PhysiCell. TiSim used an implicit-time Crank-Nicholson scheme.

### Concentration profiles in a box with one cell as point source

The following equation models the diffusion of a compound with a single point source uptake (by a cell) at a specific location in the box.

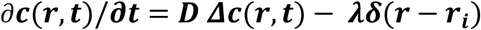

Where :

i : cell index

r_i : cell position

***c***: concentration in *μμ*mol/l or *μμ*M D: diffusion coefficient in um²/min

*λ* : uptake/decay rate in mol/min per cell

In this test, we assume a box of 60 μm each side with a single cell located at the centre of the box that acts as a point sink (see Figure 2A). The cell remains static and inert throughout the simulation, devoid of growth or movement. Dirichlet boundary conditions at the six box walls are set to 10 μM. We assume the box is empty initially. The cell metabolizes 1.6E-16 mol/min. The time resolution is set to 0.01 minutes and the spatial resolution is set to 20 μm.

**Figure 1:**
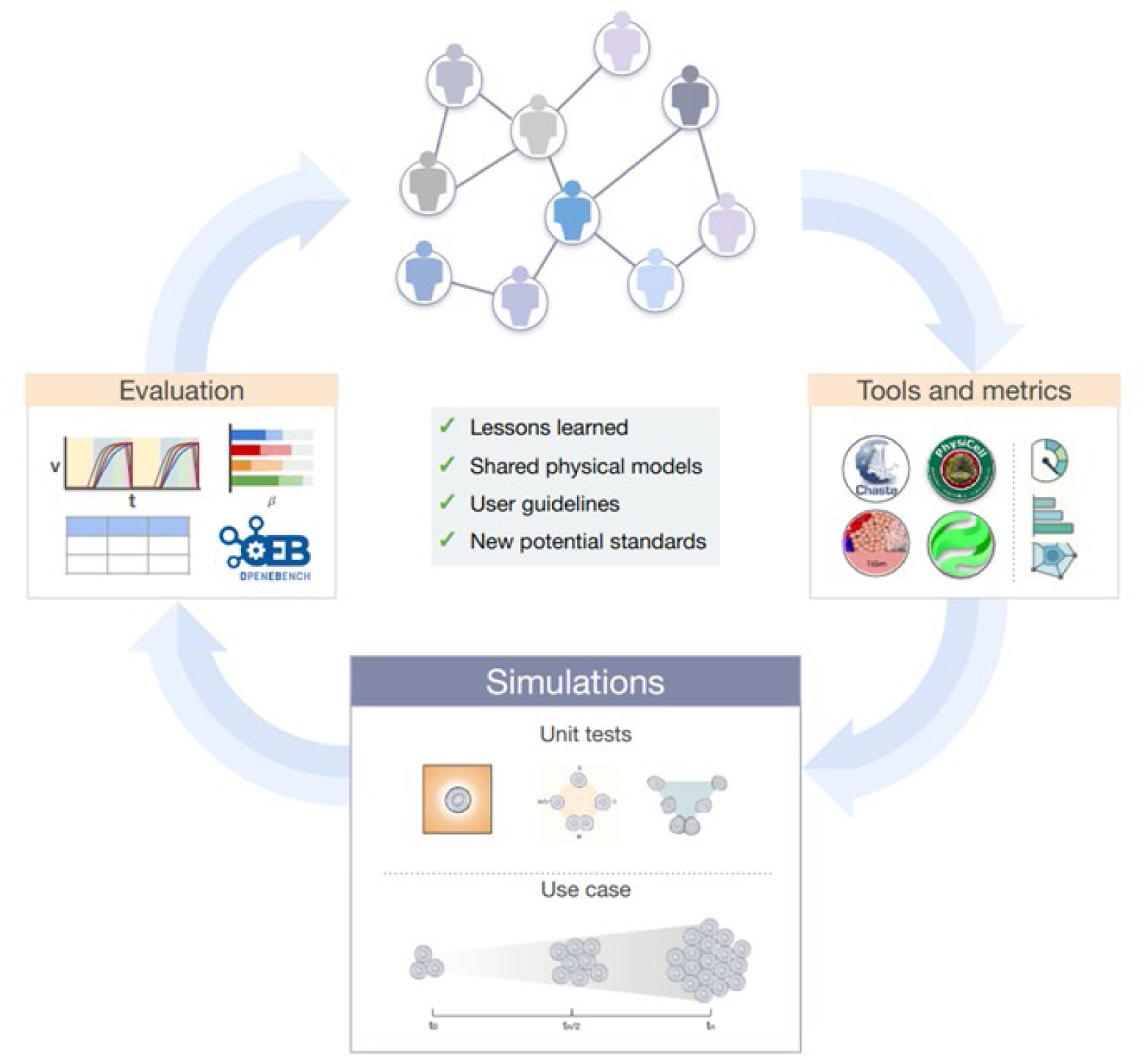
Overview of the community benchmark of CBM tools. After a call to the community, researchers from 4 tools gathered and devised tests to compare their simulation tools. The tests focused on key modelling functionalities common to all tools and that could be used as building blocks of bigger simulations in cell biology. These tests were then run in all the tools and their results were evaluated and disseminated in publicly available websites.

**Figure 2:**
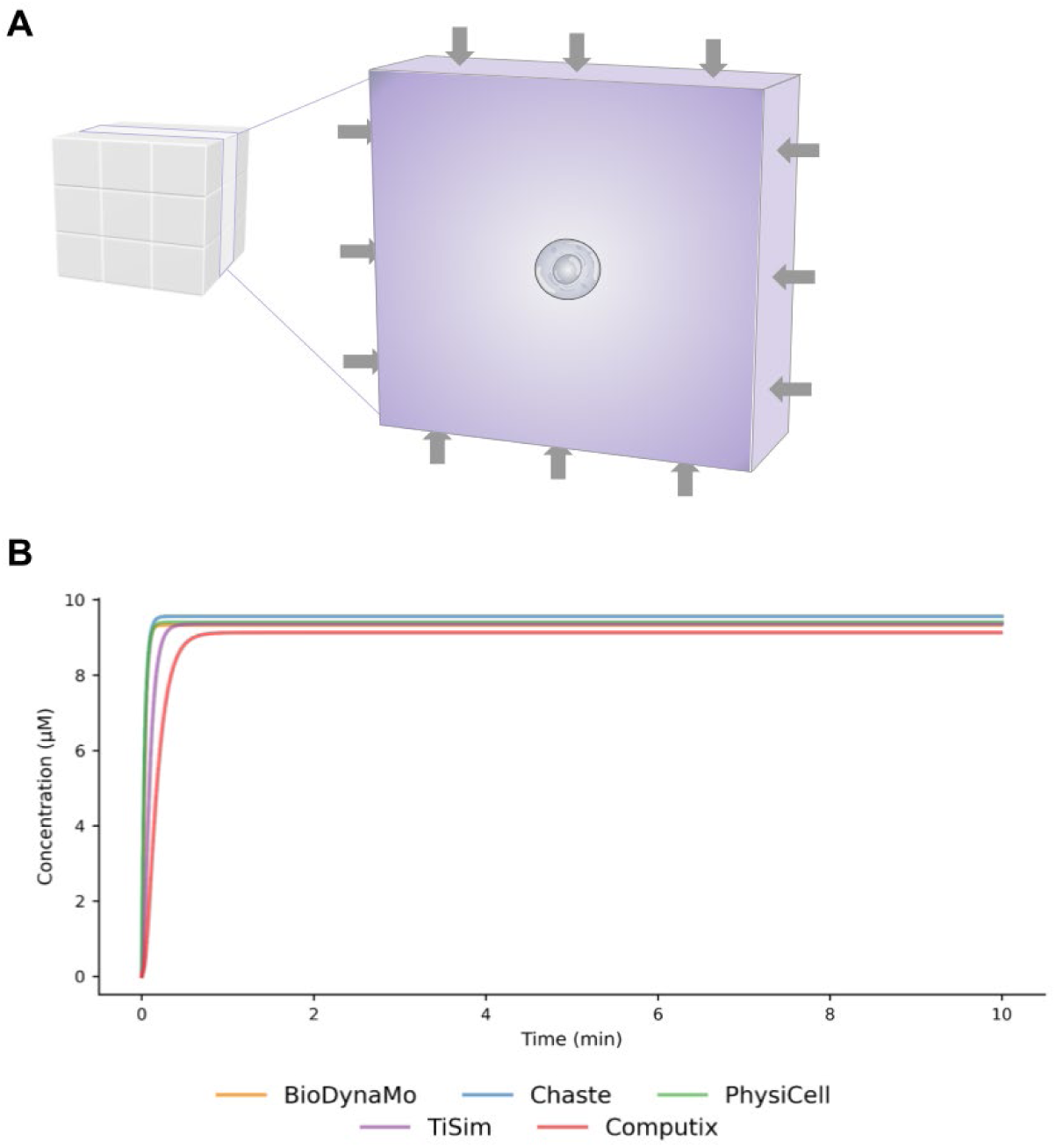
The diffusion unit test across the multiscale tools. A) Schema of the unit test where the substance is being produced at the boundaries and one cell sink sitting in the center uptakes substrate. B) Simulated concentration of a substance in the midpoint of the domain over time.

The simulation is run until a steady state is reached, which as the dynamics of this test are quite fast, occurs after a short time (Figure 2B). Looking at the concentration in the midpoint of the domain, the mean equilibrium of the tools are 9.415 μM, with Chaste showing the greater 9.56, CompuTiX the lower 9.13 and PhysiCell (9.41), TiSim (9.36) and BioDynaMo (9.33) in between.

This test is foremost used as a qualitative comparison of the tools and their capacities. We are currently working on extending this use case to pinpoint commonalities and differences across the tools and to quantify the variations in results in relation to the methods, approaches and implementations that are used. In general, the differences observed in Figure 2B could be interpreted as caused by the different numerical schemes used by each tool to solve the same system of equations. Also, as negative concentrations are not physically meaningful, they were avoided either by multiplying *λλ* by a Heaviside function or by avoiding removal of more molecules than present in a specific voxel, causing that the steady state concentration of the tools can vary.

### Unit tests for individual cell and cell-to-cell mechanical interactions

The purpose of the biomechanical unit tests is to evaluate the contact models, their numerical solutions, and their impact on the simulation results for when two biological cells get in close proximity, contact or overlap.

All CBM tools tested include models for point fixed-body mechanics and can facilitate tracking the collisions of cells amongst them and with different objects in space. Each tool includes one or several contact models, such as Hertzian contact with no adhesive forces (Mindlin & Deresiewicz, 1953), extended Hertz with an adhesion force using the Hertz contact area, used by Chaste, PhysiCell and TiSim, Johnson-Kendall-Roberts (JKR) (Chu et al., 2005; Johnson et al., 1971) with an adhesion force using the real contact area used by CompuTiX, or a phenomenological model (Zubler & Douglas, 2009), used by BioDynaMo (see Supplementary File 1 for more details on the equations used by the models).

The set-up of this test was having cells being pushed by forces and studying their reaction to these. We varied the tests to study the movement of a cell and the relaxation of cells when in contact. We designed 2 unit tests: the first one involves a single cell being pushed by applying an instantaneous concentrated force, while the second test considers two cells approaching each other, they approach, overlap and relax to an equilibrium distance.

### Studying the movement of a cell after a single push

In this simple test, we verify the dynamics of a single cell by subjecting it to a given force. This test basically mimics the active movement of a cell in a homogeneous and isotropic friction environment, the extracellular matrix (ECM).

A single cell of 10 μm diameter is situated within a box of 60 μm-wide sides. In the first timestep the cell has a force applied such that it generates movement at a speed of 10 μm/s (Figure 3A). The time resolution is 0.1 min, and the duration of the simulation is until the dynamic reaches a stable state.

**Figure 3:**
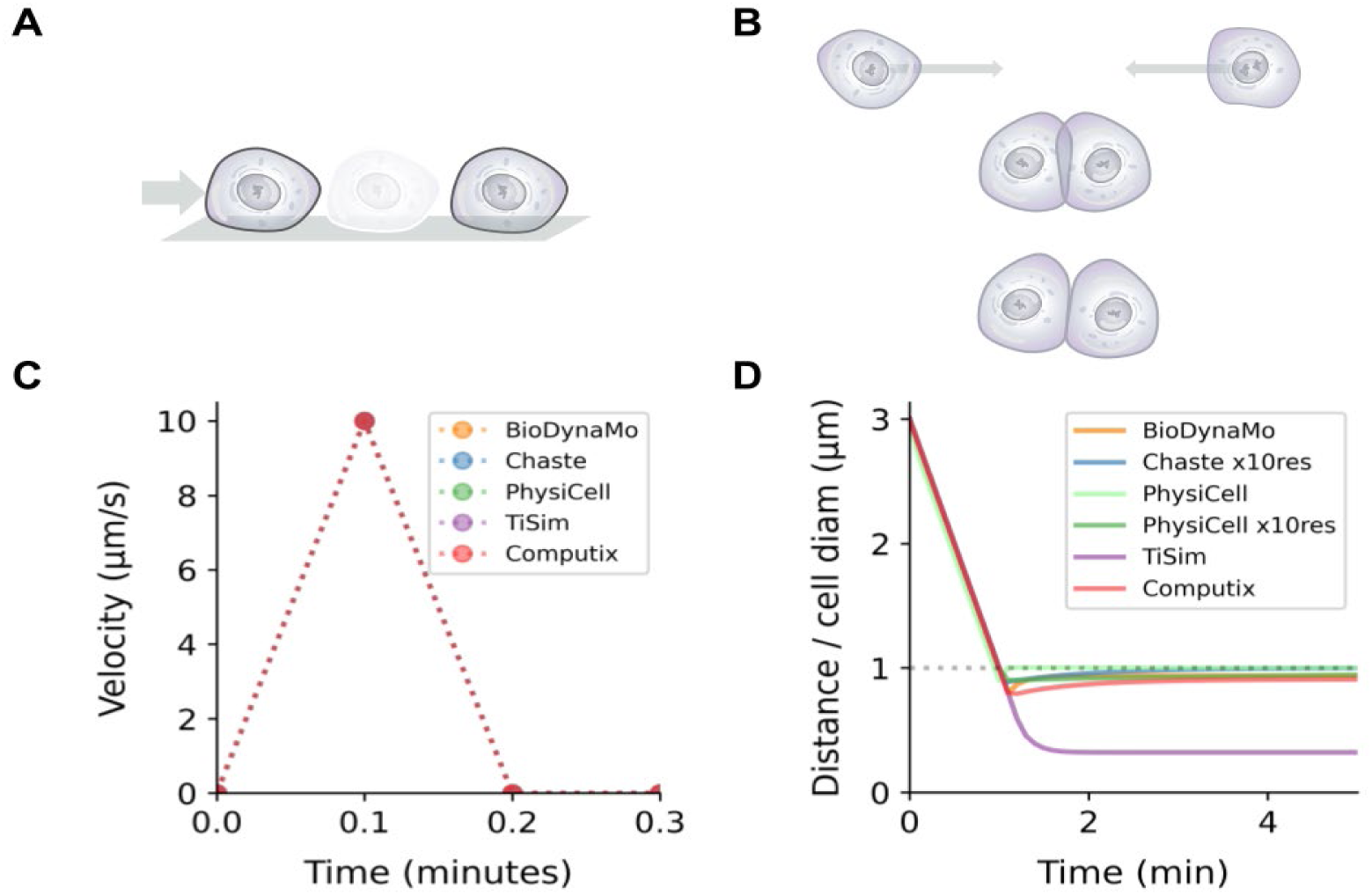
The mechanics unit tests across the multiscale tools. A) Schema of the unit test of the movement of a cell after a single push. B) Schema of the unit test of two cells approaching and repelling after their overlap. C) Dynamics of the velocity of one cell after a single push. D) Dynamics of the relative distance between cells when approaching. For TiSim, the external force was not removed (see text).

As all tools use non-inertial movement, the velocity is calculated by the following relation:

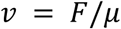

Where *F* is the force applied to the cell and *μμ* is a friction coefficient. The results showed that the cell had an instantaneous velocity of 10 um/s and then stopped in all of the tools (Figure 3C). The velocity of the cell is then integrated to obtain its position using an Euler scheme. Despite the simplicity of the test, this is a necessary step in verifying the force models across different computational codes.

### Studying the relaxation of two cells after contact

Next we aimed to evaluate how the different tools deal with the mechanics during contact between cells by simulating the mechanical relaxation of two cells when in contact. We study the effect of the different contact models used by the tools during this situation.

The set-up of this test consists of an initial condition where two cells are separated by 30 μm of their center of mass. Then, an external force is applied so that the cells move with a speed of 10 μm/min towards each other until they touch each other. The force is maintained until the overlapping cells have a distance of 0.9 x diameter, meaning they have 10% overlap. Finally, once the cells reach this distance threshold, the external forces disappear and the cells move away from each other as a consequence of the repulsion force resulting from their overlap (Figure 3B).

Figure 3D shows how the different force-based contact models implemented by the tools affect the distance between the cells’ centre as a function of time. The tools relax the contact when the cells have a 10% overlap and reach a relaxation distance that varies a bit among the tools. We chose 10% diameter overlap as higher values using different contact models cause big differences in the repulsion forces of the different contact models (Van Liedekerke et al., 2019).

BioDynaMo, CompuTiX, Chaste and PhysiCell cells overlap their volumes and then relax to the diameter distance due to the repulsive forces. In TiSim the external force applied on the cell has not been removed so it reached a maximum overlap of 0.62 x diameter.

### Unit tests for the cell cycle progression

The purpose of the cell cycle unit test is to evaluate the volume change of cells upon growth, as well as their ability to reproduce a given growth dynamic and to reproduce the four phases of a cell cycle model (Figure 4A).

**Figure 4:**
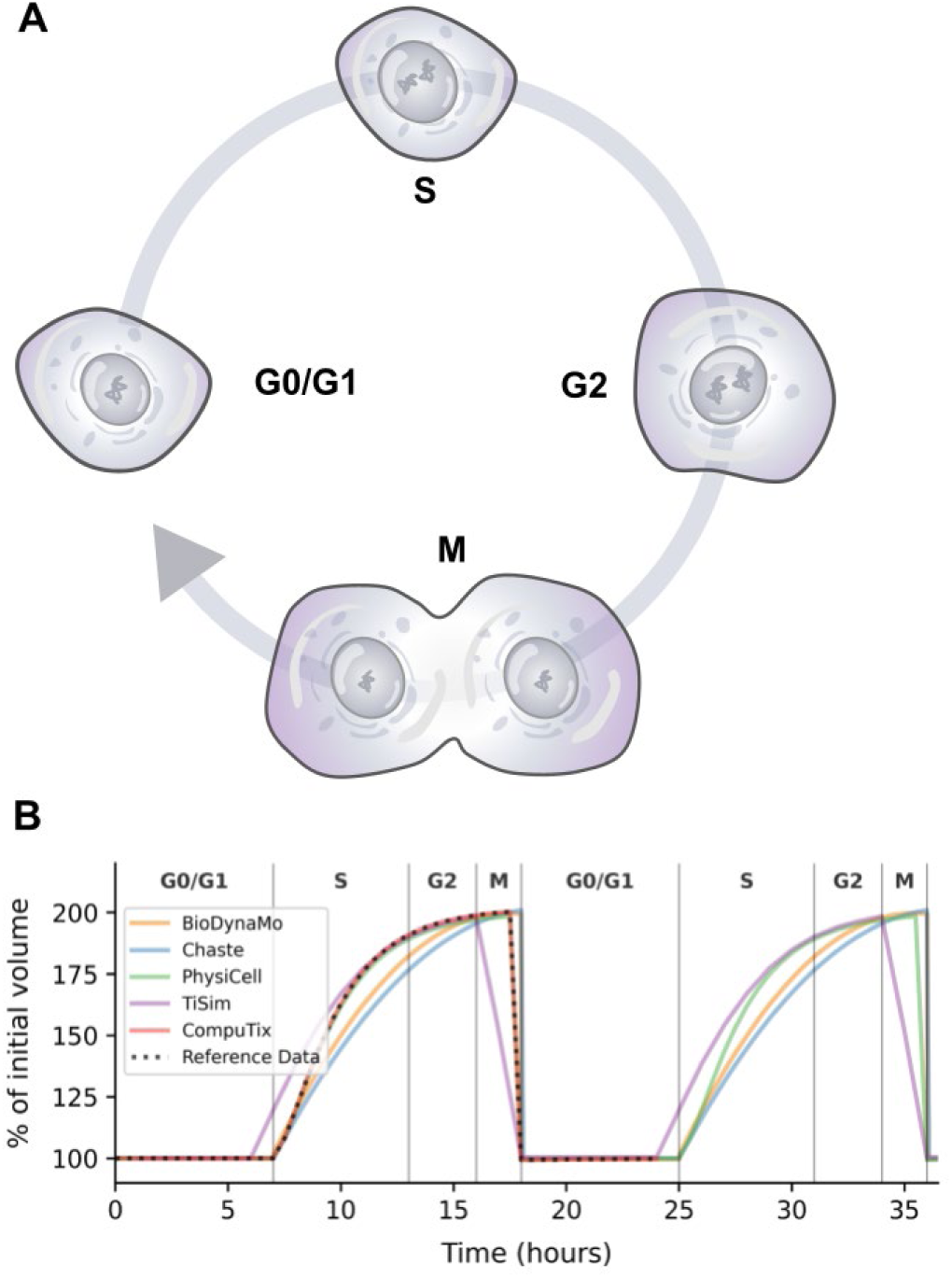
The cell cycle unit tests across the multiscale tools. A) Schema of the 4-phases cell cycle used in this test. B) Dynamics of cell volume change with fixed durations.

All the tested CBM tools have implementations to model cell volume growth, cell cycle progression, and cell division. These cell cycles are highly customizable in all tools and can showcase a one-stage cycle with a constant division rate, or a two-stage cell cycle, or even a four-stage cell cycle, with and without checkpoints, etc. The progression of the cells in these cycles can have discrete fix-time phases, can be ODE-based or rule-based and can show deterministic or stochastic transitions.

The set-up of this test consisted in following the volume growth of cells undergoing cell cycles using fixed phases duration. To make the simulations physiologically relevant, we aimed at replicating the “separated flow cytometry” 4-phases cell cycle model from the MultiCellDS work (Friedman et al., 2016) (Figure 4A) and follow a logistic growth dynamic (see Methods).

### Studying a cell cycle of fixed duration

The goal of this test was to compare how different tools handle cell cycles with a fixed duration by inspecting the volume dynamics of a single cell. We devised an initial single spherical cell, not moving, in a box of 60 μm side, that follows a cell cycle with four phases with fixed durations: G0/G1 takes 7 hours, S takes 6 hours, G2 takes 3 hours, and M takes 2 hours. The total is 18 hours for a complete growth and division cycle. The time resolution is set to 6 minutes, and the duration of the simulation is 48 hours.

The tools show qualitatively similar dynamics among them and to the reference data. However, TiSim manages the M phase differently by means of a transient dumbbell geometry established to avoid sudden changes of the cell shape during M-phase (Drasdo & Hoehme, 2005). The single cell growth curve has been chosen to start earlier in TiSim than in the other tools in order to keep the overall duration of the growth phase the same as in the other tools. Moreover, the rest of the tools exhibit slight differences in terms of the maximum volume attained at the end of the M phase and the growth dynamics (Figure 4B).

### Monolayer growth use case

Finally, we designed a test that would integrate a selection of the previous unit tests while testing the ability of the CBM tools to reproduce a biological experiment of a cell population growth. One of the easiest experimental set-up that these tools can simulate is the *in vitro* growth of cells on a Petri dish, where the cells grow in a monolayer such that nutrients and oxygen supply are abundant and accessible to each individual cell. Below a certain cell population size, the growth of the population is exponential, while beyond a certain monolayer diameter, the expansion of the monolayer is linear, reflecting only cells close to the monolayer border to contribute to its expansion (Drasdo & Hoehme, 2005). This effect can equally be observed in multicellular spheroids in three dimensions under nutrient-rich conditions (Hoehme & Drasdo, 2010b; Jagiella et al., 2016) indicating how important it is that the models are able to correctly reproduce mechanically forms of growth inhibition.

This use case was selected as it encompasses the interplay of functions previously evaluated in the unit tests (e.g., cell growth, mechanics and cell-cell interactions). The diffusion was not considered important as the biological experiments were done in medium-rich conditions. Additionally, having a two-dimensional setup would ease the visualisation of differences and help discerning potential underlying causes more effectively than in a three-dimensional context (e.g., tumour spheroids).

We decided to use the experiment from Bru and colleagues (Brú et al., 1998) that was also analysed in other works (Drasdo & Hoehme, 2005) where they used a nutrient-rich media on Petri dishes and tracked the perimeter and diameter of the cells’ monolayer. Unlike previous unit tests, where we relied upon analytical solutions or desired behaviours, in this use case the experimental results were used to benchmark the accuracy of the CBMs simulations.

The experimental data for the monolayer indicated a cell population disk diameter that ranged from 1140 μm on the first time point (day 14) to 3040 μm on the fifth and last time point (day 27). For each tool, the goal was to implement the CBM simulation that would produce results fitting those data points. Most of the tools generated results that qualitatively fit the data points well. However, the rate by which the tumour developed showed slight differences across tools (Figure 5A) as well as the deviation from the experimental data (Figure 5B).

**Figure 5.**
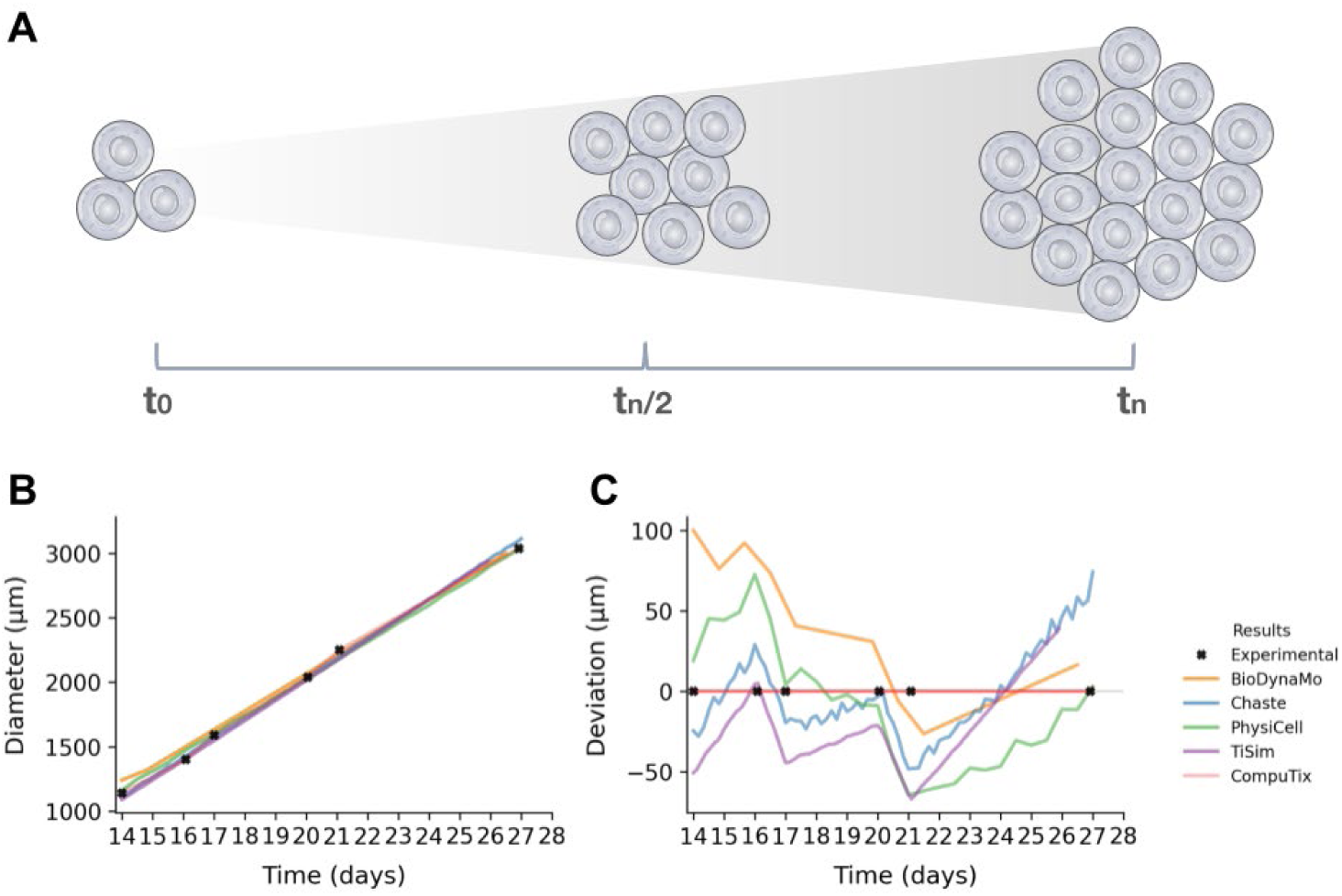
The monolayer of cells use case across the multiscale tools. A) Schema of a monolayer of cells growing. B-C) Dynamics of a monolayer of cells growing in a Petri dish. Monolayer diameter (B) and deviation from the linear interpolation of the experimental data (C) changes between days 14 and 27. Experimental data from (Brú et al., 1998).

### Disseminating the results of this benchmark

Sharing the results of this benchmark study is of great value to the modelling community in agent-based modelling and is anticipated to promote broadly progress in computational biology, computer science and technology. Furthermore, we invite members of this wider community to actively participate in this initiative. To support this, we have openly shared our processes, methodologies, tests, code, and results, enabling others to reproduce and enhance this benchmark. Through this, we aim to foster trust and ensure that our findings are robust, reliable, and impactful for the whole computational biology modelling field.

We have set up a public platform, in collaboration with OpenEBench (Capella-Gutierrez et al., 2017), to present the benchmarks’ results and methodologies, serving as reference for teams developing future simulation tools. OpenEBench is a benchmarking and technical monitoring platform that provides quality metrics for life sciences tools and workflows. It is a resource to find, deploy, and benchmark software tools in bioinformatics. Our benchmark can be accessed at OpenEBench (https://openebench.bsc.es/projects/OEBC009) and at Zenodo (https://doi.org/10.5281/zenodo.15911390).

Additionally, we have used parts of the ELIXIR Tools Platform, specifically the *bio.tools* repository (Ison et al., 2019), to ensure that the community will have access to the results generated and comprehensive information on the tools used. The *bio.tools* is a comprehensive registry for bioinformatics resources that aims to assist researchers in various life science disciplines in identifying and utilising necessary resources and tools. The tools used in the present benchmark are freely available in the PerMedCoE catalogue of *bio.tools*: https://permedcoe.bio.tools.

Lastly, we have hosted all running available codes, methodologies, reference datasets, results files and scripts used to build the figures of present paper in this GitHub repository: https://github.com/PerMedCoE/observatory_benchmark.

## Discussion

### Comparative analysis reveals consistency among CBM simulation tools in core functionalities, with divergences arising from numerical methods and model assumptions

To deliver the benefits of Virtual Human Twins (VHTs) in biomedical research, we consider of fundamental importance to systematically compare and comprehensively evaluate different CBM tools that have proven their usefulness in personalised medicine projects. To reach this goal, this contribution presents a community-based benchmark exercise carried out by five groups that are core developers of CBM simulations tools (i.e., BioDynaMo, CompuTiX, Chaste, PhysiCell, TiSim) with the aim of developing the first set of evaluation standards, tools and practices. This work represents a first step towards the sustained community effort assessing existing and prospective simulation tools, while also defining a gold standard for the benchmark of these and building collaborative tools, with the final aim of enabling the use of simulations as VHTs.

We hereby report the results of the developers’ agreement on a set of tests that capture the CBM tools’ capabilities in a faithful manner, define the scope, metrics, and reference datasets of the benchmark, and provide detailed descriptions of these in publicly available sites, to facilitate comparison with other tools. Notably, this initial implementation will need to naturally evolve beyond this work and adapt to a changing research landscape, as it is happening in other communities, such as CASP with the advent of AlphaFold (Jumper et al., 2021; Kryshtafovych et al., 2023).

The proposed tests were selected to capture the most characteristic, minimal function signatures of the multiscale nature of a CBM: diffusion of substances, mechanics models and cell cycle models. For the mechanics unit tests, we report an exceptional agreement among tools in the one-cell unit test and differences in the two-cell unit test originated by the different force-based contact models implemented by the tools. This is caused by the fact that some tools use extended Hertzian contact models while others use JKR or phenomenological contact models, causing small differences when studying the relaxation forces of the cells overlapping, even though the results are still consistent with the underlying physics. Note that some tools provide the user with different contact models, so it’s up to them to choose which model better fits their simulation.

For the diffusion unit test, slight differences were observed on the final steady state reached by the tools as well as in their dynamics. We consider that these small differences are due to the different numerical method used to discretise and integrate the diffusion-reaction equation behind this unit test.

For the cell cycle unit test, the tools mostly agree on the fixed-time unit test, with slight differences on the volume growth’s dynamics. CompuTiX and TiSim are the most different tools in this regard due to their ability to simulate the dumbbell division of the cells starting M phase to maintain cell-to-cell contacts upon cell division (Drasdo & Hoehme, 2005).

Finally, we compared the tools by simulating a use case that involved the combination of the mechanics and volume growth tests and increased the complexity and scale of the simulations. For this monolayer use case, we could leverage experimental data from cells growing in vitro (Brú et al., 1998) and the outcomes of the different tools converged quite well around the experimental data. All tools implemented a form of contact inhibition, which ensured the linear expansion of the monolayer radius above a certain size.

All in all, the tools showed very similar results for most of the unit tests with some differences due to the numerical implementation and/or different underlying models and equations. It is important to note that the selected tests represent a minimal set of common tests among the tools and the limitations observed in these benchmarking examples do not inherently apply to the softwares under analysis, as they usually have many more functionalities than the ones depicted in this work. This benchmark should be viewed as a comparative framework for models, enabling the evaluation of various modelling techniques and options to aid modellers, developers and users’ decision-making.

Despite notable efforts from some tools, one of the lessons learned is the need for exhaustive documentation of the diverse mathematical models and underlying assumptions inherent to these tools. This exercise would benefit the users, future tool developers and the long term re-use of the tools and their models.

### Using publicly available platforms to establish an open benchmark for computational biology simulations

We hope these results will prompt a deeper benchmarking of other tools in the future as we aim to gather attention of the multiscale simulation community on the importance of these community-driven tests. We aim that this comparison of tools leads the community in engaging in further discussion regarding the direction in which we envision the field progressing to overcome present obstacles.

Public dissemination is another lesson learned with this work. To strengthen the simulation community in the computational biology field, we have established a public website on OpenEBench (Capella-Gutierrez et al., 2017) where we have published the benchmark results and methodologies (https://openebench.bsc.es/projects/OEBC009). The aim of this platform is to act as a reference for future simulation tools, providing the possibility to contribute to this benchmark with novel tools from other researchers. Additionally, this website functions as a landing page for the bio.tools catalogue (Ison et al., 2019), thus, enabling users to easily find the tools used along with their characteristics.

### Learning from other communities point to the need for standardized interfaces and benchmarking practices

Moreover, lessons can be learned from machine learning where libraries such as Scikit-learn, TensorFlow, and PyTorch have been successful in establishing a common testbed for interfaces allowing to test different methods, tools and solutions. These tools share similar input signatures for their functions, i.e., Python is used as their primary user interface, and are easy to set up for comparison (Li et al., 2025). The community of the aforementioned libraries have also agreed on working with interfaces for common use cases, allowing users to test basic functionalities from both a technical and methodological perspective and a software and hardware perspective (Eid et al., 2025).

This level of standardisation is still missing in CBM tools. While it might be challenging to generalise CBM due to its multiscale complexity, it should still be possible to establish common function signatures, consistent I/O, and easy comparison set-ups without necessarily meaning that the tools themselves have to become simpler or cater only for simple tests. Likewise, to improve the usability and accessibility of CBM tools, the community should aim to generate function calls, problems, and interfaces that can be used by all tools. This would allow users to easily compute performance and results and modify the test accordingly, by choosing between different simulation tools depending on the specific problem and how well each tool performed on similar benchmark tasks. By learning from the successes of machine learning software developments, we aim to enhance the CBM field and make it more user-friendly and efficient.

### Community benchmarking can advance CBM tool reliability and foster cross-project alignment

This benchmark is a pivotal occasion to present to the community a set of simple unit tests used as gold standards that developers can use to compare future tools and methods. This comparison will ensure consistency and reliability of tools, facilitate reproducibility, improve code quality, accelerate debugging and support collaborative development (Daka & Fraser, 2014).

It is important to underline that this benchmark is of a technical nature and has its scope in the core components that could be considered foundational for CBMs. Hence, and in a more general sense, the cell-level simulation community (let them be tool developers, users or interested parties) needs to expand these tests to cover other aspects of the CBMs and other users’ perspectives, discuss what is considered a “good enough” result in the absence of an analytical solution, find relevant biomedical problems that can be used as use cases and reference experimental datasets, compare parameters exploration landscapes from different tools that simulate similar outcomes, etc. This work aims to be a first step for a more diverse and ambitious benchmarking that acts as the founding stone of a validated and verified VHT, such as the one envisioned in the EDITH project (Viceconti et al., 2024).

Another lesson learned of this benchmarking experience is the importance of setting interoperability standards and aligning with scientists who are actively contributing to this field in projects such as EDITH, Research Data Alliance’s Building Immune Digital Twins Working Group (https://www.rd-alliance.org/groups/building-immune-digital-twins-wg/), Artemis (https://artemis-euproject.eu/) and LISYM (https://www.lisym.org/). In this regard, the field can lean on present and past works like MultiCellDS (Friedman et al., 2016) or the EDITH standard proposal (Mayer & Golebiewski, 2024).

Furthermore, with this work we aspire to strengthen the European Open Source Cloud (EOSC) (Calatrava et al., 2023) by promoting domain-specific interoperability in line with the EOSC Interoperability Framework for multiscale modelling communities. Additionally, we are providing test cases for EOSC’s FAIR metrics, ensuring that tools meet criteria for findability and reusability and hosting benchmark datasets and workflows on EOSC via Zenodo to serve as training materials.

### Future Directions for Advancing Cell-Level Simulation Tools Toward Virtual Human Twins

In the future, we will strive to build and run performance-oriented benchmarks that will focus on assessing the computational efficiency, speed, and scalability of different cell-level simulation tools. This will provide valuable insights into the performance of these tools under different conditions and higher workloads than the ones presented here. Initiatives like the JUBE Benchmarking Environment (http://www.fz-juelich.de/jsc/jube) can facilitate the establishment of such benchmarks with HPC needs (Herten et al., 2024).

Another prospect for benchmarking concerns the integration of AI with CBM and the use of AI to validate and verify CBM simulations. The nascent field of mechanistic learning combines mechanistic modelling, like the one in CBMs, with AI methods to allow for parameter optimisation, surrogate model learning and physics-informed neural networks (Bunne et al., 2024; Metzcar et al., 2023). Additionally, there is a growing interest in testing CBM tools for their verification, validation, and uncertainty quantification (VVUQ) (Cogno et al., 2024; Richardson et al., 2020). A benchmark on the integration of these methods with CBM would ensure their consistency and reliability and allow for collaborative development of VHTs.

Lastly, we must be aware that maintaining this benchmarking platform actively requires continuous effort, even more so if we aim at evolving the tests and standards with the codebase to avoid stagnation. Funding from EU projects was used to kickstart this effort together with in kind contributions from many researchers, but a more sustainable funding source will be needed to sustain it.

Being at the dawn of VHT use in biomedicine, we hope that this benchmarking contribution will serve towards strengthening the agent-based model simulation community and will be the launchpad to address this current challenge (Fletcher & Osborne, 2022; Macklin, 2019). By providing a common framework for benchmarking and comparison, we aim to foster collaboration, driving innovation, reducing complexity, and accelerating progress in this field as CASP has done for structural biology (Mahmood, 2025). Additionally, these standards will lower the access barrier to novel, better technologies to developers and end users by allowing their use in complex workflows and benchmarks. We envision a vibrant community where researchers and developers can learn from each other, improve their tools, and advance the understanding of cell-level phenomena.

## Supporting information

Supplementary File 1: Methods and equations used by the tools in the different tests.

## Acknowledgements

The authors acknowledge the help and guidance provided by Dmitry Repchevski, José Maria Fernández and Salvador Capella-Gutierrez from the INB-ELIXIR Spanish node on the use of OpenEBench.

This work was supported by the European Commission (grants CREXDATA 101092749, PerMedCoE 951773, and EDITH-CSA 101083771, Hanami EUROHPC-JU-2022-INCO-04-01), the UK BBSRC (Biotechnology and Biological Sciences Research Council) grant numbers BB/V018930/1, BB/V01840X/1, BB/V018647/1.

JMO’s research is supported by the ARC (Australian Research Council) grant numbers DP230100380, FT230100352. AM acknowledges funding from the Generalitat Valenciana’s CIDE GenT programme under the project CIDEXG/2023/22. MR acknowledges funding from the Ayuda Severo Ochoa CEX2021-001148-S from MICIU/AEI.

DD, JZ, JD, acknowledge funding by the BMBF (German federal ministry of Education and Research) projects LiSyM-Cancer and LiSyM-Cancer II. DD, MP, JD, JP and PVL acknowledge funding from European grant EDITH. DD, JP and MP acknowledge funding by European grant ARTEMIS.

## Supplementary materials

**Table S1.**
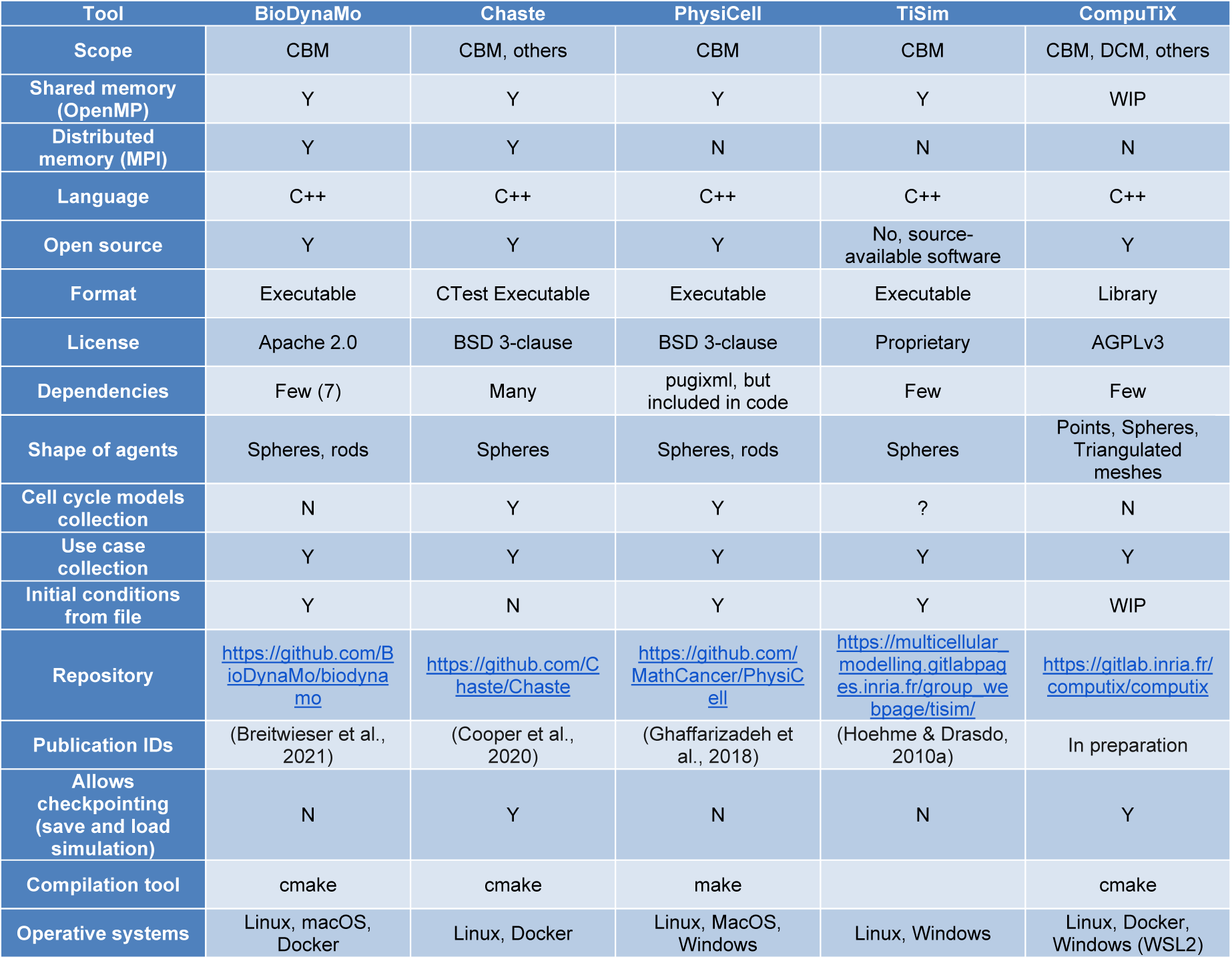
Characteristics of tools for cell population modelling contacted to benchmark. CBM stands for centre-based models. For entries represented by question marks, no evidence was found to the best of our efforts.

