## Supplementary File 1: Methods and equations used by the tools in the different tests. for "Open Benchmarking for Cell-Based Multiscale Models: Lessons from a Community Initiative"

July 16, 2025

### 1 Methods and equations used by the tools in the different tests

#### 1.1 BioDynaMo

Information can be found in these papers [Breitwieser et al., 2021].

##### 1.1.1 Diffusion

**DiffuseWithDirichlet(dt)**

$$c2[i,j,k] = c1[i,j,k]\,(1-\mu\cdot\Delta t) + \left(D\cdot\Delta t\cdot\frac{1}{\Delta x^2}\right)(c1[i-1,j,k]+c1[i+1,j,k] \\ +c1[i,j+1,k]+c1[i,j-1,k]+c1[i,j,k-1]-6\cdot c1[i,j,k]+c1[i,j,k+1]) \tag{1}$$

**Where:**

- **c1(i,j,k)**: concentration at time step t of coordinates (i,j,k).
- **c2(i,j,k)**: concentration at the time step (t+0.5) of coordinates (i,j,k).
- $\mu$: decay rate
- **D**: diffusion coefficient
- **Δx**: voxel size length
- **Δt**: time step (unit case value = 0.01)

**DiffusionGrid::ChangeConcentrationBy Parameters**

- **dx**: position of the grid (0,0,0).
- **amount**: value to change concentration (-sink*simulation_time_step).
- **mode**: change mode: additive (default), exponential, logistic
- **scale_with_resolution**: default false

**Additive**

$$c3(i,j,k) = c2(i,j,k) - S\cdot\Delta t \tag{2}$$

**Exponential**

$$c3(i,j,k) = c2(i,j,k)\cdot S\cdot\Delta t \tag{3}$$

**Logistic**

$$c3(i,j,k) = c2(i,j,k) + \begin{cases}(upper\_bound - c2(i,j,k))\cdot(-S\cdot\Delta t) & if - S\cdot\Delta t > 0\\ (c2(i,j,k) - lower\_bound)\cdot(-S\cdot\Delta t) & if - S\cdot\Delta t <= 0\end{cases} \tag{4}$$

**Where:**

- **c2(i,j,k)**: concentration at time step (t+0.5) of coordinates (i,j,k).
- **c3(i,j,k)**: concentration at the time step (t+1) of coordinates (i,j,k).
- **S**: sink rate (unit case value = 20)
- **Δt**: time step (unit case value = 0.01)

#### 1.1.2 Mechanics

BioDynaMo uses a phenomenological model in accordance with the Cortex3D implementation [Zubler and Douglas, 2009], comprising a repulsive and an attractive component

**Mechanical force exerted by a neighbor cell (cell 2) on a reference cell (cell 1)**

$$F_{21} = k * \delta - \gamma * \sqrt{r * \delta}$$

Where:

$$\begin{aligned} k &= \text{repulsion coeff} \\ \gamma &= \text{attraction coeff} \\ r &= \frac{r_1 * r_2}{r_1 + r_2} \\ \delta &= \max(0, r_1 + r_2 - r_{12}) \\ r_{12} &= \text{distance between cell centers} \\ r_1 &= \text{reference cell radius} + r_{\text{add}} \\ r_2 &= \text{neighbor cell radius} + r_{\text{add}} \\ r_{\text{add}} &= 10 * \min(\text{ref iof coeff}, \text{neigh iof coeff}) \\ \text{ref iof coeff} &= \text{neigh iof coeff} = 0.15 \end{aligned}$$

Where "iof coeff" is the Inter-Object Force coefficient and allows for effective radius different than the geometrical radius of the spherical object.

**Custom.h: dissipative force equation**

$$\vec{v_2} = \vec{v_1} - \vec{fc} \cdot \Delta t \tag{5}$$

**Where:**

- $\vec{v_1}$: velocity vector at time step (t).
- $\vec{v_2}$: velocity vector at time step (t+1).
- $\vec{fc}$: friction coefficient (value = 0.1).
- Δt: time step (value 1 equivalent to 0.1 min)

**Custom.h: apply force**

$$\vec{pos}_2 = \vec{pos}_1 + \vec{V_0} \cdot \Delta t \tag{6}$$

**Where:**

- $\vec{pos}_1$: position vector at time step (t).
- $\vec{pos}_2$: velocity vector at time step (t+1).
- $\vec{V_0}$: initial velocity vector (10,0,0).
- Δt: time step (value 1 equivalent to 0.1 min)

#### 1.1.3 Cell cycle

Biodynamo models the cell cycle using cumulative probability to determine phase progression. The time a cell has spent in its current phase is divided by the phase's expected duration, and this ratio is compared to a random number sampled from a uniform distribution. If the ratio exceeds the random number, the cell advances to the next phase.

### 1.2 Chaste

More information of equations and simulation methods can be found in [Osborne et al., 2017] and [Cooper et al., 2020].

#### 1.2.1 Diffusion

We consider diffusion of a species $u$ with cells as sinks/ sources:

$$\frac{\partial u}{\partial t} = \nabla \cdot (D \nabla u) + (k_1 u + k_2)\rho(\mathbf{x}) + (k_3 u + k_4). \tag{7}$$

Where $\rho(\mathbf{x}) = \sum_{\text{Cells}} \delta((x) - (x)_i)$ is calculated by summing the number of cells in the element scaled by area of element (solved using a Finite Element Method).

**Where:**

- $D$ is the diffusion coefficient.
- $k_1, k_2$ are the linear and constant cell dependent production and removal.
- $k_3, k_4$ are the linear and constant cell independent dependent production and removal.

#### 1.2.2 Mechanics

Following [Osborne et al., 2017] The mechanics can be described as follows: cells are represented by their centres, which are modelled as a set of points $\{\mathbf{r}_1, \ldots, \mathbf{r}_{N_{\text{Cells}}}\}$ which are free to move in space. The first-order equation of motion for each cell centre, $\mathbf{r}_i$ is given by

$$\eta \frac{\mathrm{d}\mathbf{r}_i}{\mathrm{d}t} = \sum_{j \in \mathcal{N}_i(t)} \mathbf{F}_{ij}(t), \tag{8}$$

where $\eta$ denotes a damping constant and the total force acting on a cell $i$ at time $t$ is assumed to equal the sum of all forces, coming from the connections with all neighbouring cells $j \in \mathcal{N}_i(t)$ adjacent to $i$ at that time, $\mathbf{F}_{ij}(t)$.

We define the force between cells $\mathbf{F}_{ij}(t)$ as [Osborne et al., 2017]

$$\mathbf{F}_{ij}(t) = \begin{cases} \mu_{ij} s_{ij}(t) \hat{\mathbf{r}}_{ij}(t) \log\left(1 + \frac{||\mathbf{r}_{ij}(t)|| - s_{ij}(t)}{s_{ij}(t)}\right), & \text{for} \quad ||\mathbf{r}_{ij}(t)|| < s_{ij}(t), \\ \mu_{ij}(||\mathbf{r}_{ij}(t)|| - s_{ij}(t)) \hat{\mathbf{r}}_{ij}(t) \exp\left(-k_C \frac{||\mathbf{r}_{ij}(t)|| - s_{ij}(t)}{s_{ij}(t)}\right), & \\ \qquad \text{for} \quad s_{ij}(t) \leq ||\mathbf{r}_{ij}(t)|| \leq r_{\max}, & \\ 0, \quad \text{for} \quad ||\mathbf{r}_{ij}(t)|| > r_{\max}. \end{cases} \tag{9}$$

Here $\mu_{ij}$ is known as the “spring constant” and controls the size of the force (and depends on the cell types of the connected cells), by default $\mu_{ij} = \mu$ for all interactions, $\mathbf{r}_{ij}(t) = \mathbf{r}_i(t) - \mathbf{r}_j(t)$, $\hat{\mathbf{r}}_{ij}(t)$ is the corresponding unit vector, $k_C$ is a parameter which defines decay of the attractive force, and $s_{ij}(t)$ is the natural separation between these two cells. Here $s_{ij}(t)$ is the sum of the radii of the two cells, and a cell's radius increases from 0.25 to 0.5 CDs over the first hour of the cell cycle, and hence is a function of time. Note that there is a cut off distance, $r_{\max}$, such that once $||\mathbf{r}_{ij}(t)|| > r_{\max}$ the cells are not connected so the force is zero.

#### 1.2.3 Cell cycle

Cells divide as detailed in each example. As in [Osborne et al., 2017] on cell division, we generate a random mitotic unit vector $\hat{\mathbf{m}}$ and the daughter cells are placed at $\mathbf{r}_i \pm \epsilon \hat{\mathbf{m}}$, where $\epsilon$ is a constant division separation parameter.

### 1.3 Computix

#### 1.3.1 Diffusion with cells in the center

**Problem statement**

This simulation illustrates diffusion of a species in a box with multiple point sinks representing cells with a constant uptake rate. The equation to solve is then a reaction-diffusion equation with a homogeneous source term.

$$\partial_t c(\vec{x},t) - \nabla \cdot (D \nabla c(\vec{x},t)) = -\sum_l \lambda_l \delta_{\vec{x}_l}(\vec{x}) \qquad \text{if } \vec{x} \in \left(-\frac{L}{2}, \frac{L}{2}\right)^3$$
$$c(\vec{x}) = c_{\text{BC}} \qquad \text{if } x,y,z = \pm\frac{L}{2}$$
$$c(\vec{x},0) = c_{\text{BC}}$$

With $c$ the concentration, $c_{\text{BC}}$ the initial concentration and also concentration at the boundary, $D$ the diffusion coefficient and $\lambda_l$ the uptake rate of the cell $l$, $\delta_{\vec{x}_l}$ the Dirac's function at the cell's position $\vec{x}_l$ and $L$ the length of the box.

**Analytical solution**
The analytical solution is given as

$$c(\vec{x},t) = c_{\text{BC}} - \sum_l \lambda_l \sum_{ijk}^{\infty} \phi_{ijk}(\vec{x})\phi_{ijk}(\vec{x}_l) \frac{1 - \mathrm{e}^{-\lambda_{ijk} t}}{\lambda_{ijk}}$$
$$\phi_{ijk}(x,y,z) = \left(\frac{2}{L}\right)^{\frac{3}{2}} \cos\left(\frac{(2i+1)\pi x}{L}\right) \cos\left(\frac{(2j+1)\pi y}{L}\right) \cos\left(\frac{(2k+1)\pi z}{L}\right)$$
$$\lambda_{ijk} = \left((2i+1)^2 + (2j+1)^2 + (2k+1)^2\right) \frac{\pi^2 D}{L^2}$$

**Finite Volume scheme**
The physical domain is a cube of length $L$ and volume $V = L \times L \times L$. The domain is then discretized regularly with cubes of length $a$.

The equation is solved numerically using a finite volume approach which is described here after. For a voxel $\alpha$, integrating on its volume $V_\alpha$.

$$\partial_t \int_{V_\alpha} \mathrm{d}V\, c = \int_{V_\alpha} \mathrm{d}V\, D\nabla^2 c - \int_{V_\alpha} \mathrm{d}V \sum_{l:\vec{x}_l \in V_\alpha} \lambda_l \delta_{\vec{x}_l}(\vec{x}) \tag{10}$$

which gives, considering $c_\alpha = \frac{n_\alpha}{V_\alpha}$, Gauss's theorem, and the integration of Dirac delta functions,

$$\partial_t n_\alpha = \oint_{\partial V_\alpha} \mathrm{d}\vec{A} \cdot D\nabla c - \sum_{l:\vec{x}_l \in V_\alpha} \lambda_l \tag{11}$$

Then,

$$\partial_t n_\alpha = \sum_{\beta \in \text{Neigh}(\alpha)} A_{\alpha\beta} D \frac{c_\beta - c_\alpha}{d_{\alpha\beta}} - \sum_{l:\vec{x}_l \in V_\alpha} \lambda_l$$
$$= \frac{D}{a^2} \sum_{\beta \in \text{Neigh}(\alpha)} (n_\beta - n_\alpha) - \sum_{l:\vec{x}_l \in V_\alpha} \lambda_l$$

where the first sum is over the neighbouring voxels of $\alpha$, and the second is over all cells that belong to voxel $\alpha$. We assume $d_{\alpha\beta} = a$, the distance between the centers of voxels $\alpha$ and $\beta$.

The implicit scheme is given by:

$$n_\alpha^{t+\Delta t} - C_{\text{Courant}} \sum_{\beta \in \text{Neigh}(\alpha)} \left(n_\beta^{t+\Delta t} - n_\alpha^{t+\Delta t}\right) = n_\alpha^t - \Delta t \sum_{l:\vec{x}_l \in V_\alpha} \lambda_l \tag{12}$$

with $C_{\text{Courant}} = \frac{D\,\Delta t}{a^2}$ the Courant number. This linear system can then be solved with a conjugate gradient solver.

Dirichlet boundary conditions are set by introducing ghost voxels (see figure below), with the same volumes as internal ones and surrounding the box. These ghost voxels have constant concentration $c_{\text{BC}}$.

#### 1.3.2 Diffusion and decay

**Problem statement**
This simulation illustrates the diffusion of a species in a box with a homogeneous uptake rate everywhere in the domain. The equation to solve is a reaction-diffusion equation with a homogeneous source term:

$$\begin{aligned}
\partial_t c(\vec{x},t) - \nabla \cdot (D\nabla c(\vec{x},t)) &= -\lambda c(\vec{x},t) && \text{if } \vec{x} \in \left(-\frac{L}{2}, \frac{L}{2}\right)^3 \\
c(\vec{x}) &= c_{\mathrm{BC}} && \text{if } x,y,z = \pm\frac{L}{2} \\
c(\vec{x},0) &= 0
\end{aligned}$$

Here, $c$ is the concentration, $c_{\mathrm{BC}}$ is the boundary and initial concentration, $D$ is the diffusion coefficient, $\lambda$ is the uptake rate, and $L$ is the length of the box.

**Analytical Solution**

The analytical solution is given by:

$$\begin{aligned}
c(\vec{x},t) &= c_{\mathrm{BC}} - \lambda c_{\mathrm{BC}} \sum_{ijk}^{\infty} \phi_{ijk}(\vec{x}) \mathrm{e}^{-\lambda_{ijk} t} \int_0^t \int_\Omega \mathrm{d}t_0 \, \mathrm{d}V(\vec{x}_0) \, \phi_{ijk}(\vec{x}_0) \mathrm{e}^{\lambda_{ijk} t_0} \\
&= c_{\mathrm{BC}} - \lambda c_{\mathrm{BC}} \sum_{ijk}^{\infty} A_{ijk} \phi_{ijk}(\vec{x}) \frac{1 - \mathrm{e}^{-\lambda_{ijk} t}}{\lambda_{ijk}} \\
A_{ijk} &= \frac{8L^3}{\pi^3 (2i+1)(2j+1)(2k+1)} \left(\frac{2}{L}\right)^{3/2} \\
&\quad \times \sin\left(\frac{(2i+1)\pi}{2}\right) \sin\left(\frac{(2j+1)\pi}{2}\right) \sin\left(\frac{(2k+1)\pi}{2}\right) \\
\phi_{ijk}(x,y,z) &= \left(\frac{2}{L}\right)^{3/2} \cos\left(\frac{(2i+1)\pi x}{L}\right) \cos\left(\frac{(2j+1)\pi y}{L}\right) \cos\left(\frac{(2k+1)\pi z}{L}\right) \\
\lambda_{ijk} &= \lambda + \frac{\pi^2 D}{L^2} \left[(2i+1)^2 + (2j+1)^2 + (2k+1)^2\right]
\end{aligned}$$

**Finite Volume Scheme**
The physical domain is a cube of length $L$, and volume $V = L \times L \times L$. The domain is discretized into cubes (voxels) of length $a$. We solve the equation numerically using a finite volume approach.
For a voxel $\alpha$, integrating over its volume $V_\alpha$:

$$\partial_t \int_{V_\alpha} \mathrm{d}V \, c = \int_{V_\alpha} \mathrm{d}V \, D\nabla^2 c - \int_{V_\alpha} \mathrm{d}V \, \lambda c$$

Assuming $c_\alpha = \frac{n_\alpha}{V_\alpha}$, applying Gauss's theorem, and simplifying:

$$\partial_t n_\alpha = \oint_{\partial V_\alpha} \mathrm{d}\vec{A} \cdot D\nabla c - \lambda n_\alpha$$

This gives:

$$\begin{aligned}
\partial_t n_\alpha &= \sum_{\beta \in \mathrm{Neigh}(\alpha)} A_{\alpha\beta} D \frac{c_\beta - c_\alpha}{d_{\alpha\beta}} - \lambda n_\alpha \\
&= \frac{D}{a^2} \sum_{\beta \in \mathrm{Neigh}(\alpha)} (n_\beta - n_\alpha) - \lambda n_\alpha
\end{aligned}$$

Here, the sum is over the neighboring voxels of $\alpha$, and we assume $d_{\alpha\beta} = a$, the distance between centers of voxels.

**Implicit Time Scheme**

The implicit scheme is:

$$(1+\lambda\Delta t)\, n_\alpha^{t+\Delta t} - C_{\text{Courant}} \sum_{\beta\in\text{Neigh}(\alpha)} \left(n_\beta^{t+\Delta t} - n_\alpha^{t+\Delta t}\right) = n_\alpha^t$$

with Courant number $C_{\text{Courant}} = \frac{D\Delta t}{a^2}$. This linear system can be solved using a conjugate gradient solver.

**Boundary Conditions**

Dirichlet boundary conditions are imposed using forcing voxels in the boundaries to have an assigned constant concentration $c_{\text{BC}}$.

#### 1.3.3 Movement with friction

**Problem statement**

This simulation illustrates the movement of a cell given a homogeneous isotropic friction coefficient, that mimics friction of a cell with the extracellular matrix in an idealised way, after the instantaneous application of a fixed and constant force.

**Equation of motion**
The system to solve is then composed of the two equation of motion for velocity and position:

$$\gamma\vec{v}(t) = \vec{F}_{\text{loc}}(t)$$
$$\frac{\mathrm{d}\vec{x}}{\mathrm{d}t}(t) = \vec{v}(t)$$

with $\vec{x}$ the position, $\vec{v}$ the velocity, $\vec{F}_{\text{loc}}$ the external force and $\gamma$ the friction coefficient with the external medium.

**External force** The initial constant force is applied to an initially motionless sphere, and then immediately removed:

$$F_{\text{loc}}(t) = \begin{cases} F_{\text{loc}}, & \text{if } t = 0 \\ 0, & \text{if } t > 0 \end{cases} \tag{13}$$

**Stokes drag force**
Stokes drag force is used to model the viscous friction, assuming a homogeneous isotropic friction coefficient that mimics friction of a cell with the extracellular matrix in an idealised way.

$$\vec{F}^{\text{Stokes}} = -\gamma\vec{v}^{\text{rel}} \tag{14}$$

Where $\gamma$ is the friction coefficient or friction matrix for a given particle and $\vec{v}^{rel}$ is a relative velocity of the particle with respect to the fluid.

**Integration scheme**
Set of equations we are going to integrate are defining the implicit numerical scheme.

$$M\cdot(\vec{v}(t+\Delta t) - \vec{v}(t)) = \Delta t\cdot\left[\vec{F}_{\text{loc}}(t) - \gamma\vec{v}(t)\right]$$
$$\vec{x}(t+\Delta t) - \vec{x}(t) = \Delta t\cdot\vec{v}(t+\Delta t)$$

where the mass tensor is defined as $M = m\mathbb{I} - \gamma\Delta t$. In our case we neglect the intertial constribution, $m\mathbb{I}$, resulting in an overdamped regime. One can also prove the solution for velocity is the same one would have obtained for the explicit overdamped scheme:

$$\begin{aligned} & M = \gamma\Delta t \\ & \Rightarrow \gamma\Delta t(\vec{v}_{t+\Delta t} - \vec{v}_t) = \Delta t(\vec{F} - \gamma\vec{v}_t) \\ & \Rightarrow \gamma(\vec{v}_{t+\Delta t} - \vec{v}_t) = \vec{F} - \gamma\vec{v}_t \\ & \Rightarrow \vec{v}_{t+\Delta t} = \frac{\vec{F}}{\gamma}. \end{aligned}$$

The parameters used in the use case are target time ($t_{\text{end}} = 10$ minutes), time step ($\Delta t =$ 0.1 minute), cell's radius ($r = 5$ micrometers), initial force ($F_{\text{loc}} = (\begin{pmatrix}10 & 0 & 0\end{pmatrix})$ micrometer) and friction coefficient ($\gamma = 1$ kilogram per minute).

#### 1.3.4 Pushing cells

**Problem statement** This simulation illustrates the dynamics of two cells, identical in radius and properties, pushed towards each other with an external force which is removed when their overlap reaches a critical value equal to 10% the diameter of the spheres. JKR model is tested in this example rather than the Hertz one and friction with the external medium is modeled as a viscous friction.

**Equation of motion**
The system to solve for the two spheres is then composed of the two equation of motion for velocity and position:

$$\gamma\vec{v}(t) = \vec{F}_{\mathrm{loc}} + \vec{F}_{\mathrm{Hertz}} + \vec{F}_{\mathrm{JKR}}$$
$$\frac{\mathrm{d}\vec{x}}{\mathrm{d}t}(t) = \vec{v}(t)$$

with $\vec{x}$ the position, $\vec{v}$ the velocity, $\vec{F}_{\mathrm{loc}}$ the external force, $\vec{F}_{\mathrm{JKR}}$ a collision adhesive force force and $\gamma$ the friction coefficient with the external medium.
**External force**
Two forces, equal and opposite in direction, are applied to the two spheres until the value of the intersection reaches 10% their diameter value:

$$F_{\mathrm{loc}}(\delta) = \begin{cases} F_{\mathrm{loc}}, & \text{if } \delta \leq \theta \cdot 2r \\ 0, & \text{if } \delta > \theta \cdot 2r \end{cases} \tag{15}$$

Where $\delta$ is the spheres' overlap, $r$ the radius of the two spheres and $\theta$ the threshold value, set equal to 0.1 in this case to capture the 10% overlap.
**Stokes drag force**
Stokes drag force is used to model the viscous firction, assuming a homogeneous isotropic friction coefficient that mimics friction of a cell with the extracellular matrix in an idealised way.

$$\vec{F}^{\mathrm{Stokes}} = -\gamma\vec{v}^{\mathrm{rel}} \tag{16}$$

Where $\gamma$ is the friction coefficient or friction matrix for a given particle and $\vec{v}^{rel}$ is a relative velocity of the particle with respect to the fluid.
**JKR model**
Johnson-Kendall-Roberts model, with damping proportional to contact area, can be used to model the collision of two spheres. The JKR force together with Hertz force are applied here:

$$\vec{F} = \vec{F}_{\mathrm{Hertz}} + \vec{F}_{\mathrm{JKR}} = -\frac{4}{3}E^*\frac{a^3}{R}\hat{n} + \sqrt{8\pi\gamma^* E^* a^3}\hat{n} \tag{17}$$

Where $a$ is the contact radius, $1/R$ is the curvature, $\hat{n}$ the normal direction, $E^*$ the effective Young's modulus and $\gamma^*$ the contact adhesion energy density.
**Integration scheme**
The implicit integration scheme is used and set of equations we are going to integrate is the following:

$$(\vec{v}(t+\Delta t) - \vec{v}(t)) \cdot \left[m\mathbb{I} - \Delta t\ \nabla_{\vec{v}}\vec{F}(\vec{x}(t), \vec{v}(t))\right] = \Delta t\ \vec{F}\left(\vec{x}(t), \vec{v}(t)\right)$$
$$\vec{x}(t+\Delta t) - \vec{x}(t) = \Delta t \cdot \vec{v}(t+\Delta t)$$

Where the combined mass tensor matrix contains a (negative) friction matrix term $\nabla_{\vec{v}}\vec{F}$.

$$M = m\mathbb{I} - \Delta t\ \nabla_{\vec{v}}\vec{F}(\vec{x}(t), \vec{v}(t)) \tag{18}$$

All the forces are joined as:

$$\vec{F} = F_{\mathrm{loc}} + \vec{F}^{\mathrm{Stokes}} + \vec{F}_{\mathrm{Hertz}} + \vec{F}_{\mathrm{JKR}} \tag{19}$$

Allowing for approximation of the acceleration $\vec{a}$ at time $t$ in the first equation as:

$$\vec{a}(t) = \frac{\vec{v}(t+\Delta t) - \vec{v}(t)}{\Delta t} \tag{20}$$

Can be rewritten to obtain the implicit system.

$$\vec{a}(t)M = \vec{F}(\vec{x}(t), \vec{v}(t)) \tag{21}$$

Which can be solved by using Conjugate Gradient for computing the accelerations, followed by Forward Euler for updating velocities and positions from:

$$\vec{v}(t+\Delta t) - \vec{v}(t) = \Delta t \cdot \vec{a}(t+\Delta t)$$
$$\vec{x}(t+\Delta t) - \vec{x}(t) = \Delta t \cdot \vec{v}(t+\Delta t)$$

#### 1.3.5 Cell cycle

TBD

### 1.4 PhysiCell

PhysiCell methods and equations full definition are broadly described in the Design and implementation section of [Ghaffarizadeh et al., 2018]. Additional information can be found in [Ghaffarizadeh et al., 2016].

#### 1.4.1 Diffusion Biochemical microenvironment

Reaction-diffusion PDEs with Bulk source/sinks and cell-centered sources and sinks. BioFVM uses a Cartesian mesh and an implicit-time Euler numerical scheme. The following equation is used in the diffusion unit case.

BioFVM solves PDEs of the form of:

$$\frac{\delta \vec{p}}{\delta t} = \overbrace{\vec{D}\Delta^2 \vec{p}}^{diffusion} + \overbrace{\vec{\lambda}\vec{p}}^{decay} + \overbrace{\vec{S}(\vec{p}^* - \vec{p})}^{bulk-source} + \overbrace{\vec{U}\vec{p}}^{bulk-uptake} + \overbrace{\sum_{cells\ k} 1_k(\vec{x})[\vec{S}_k(\vec{p}^* - \vec{p}) - \vec{U}_k \vec{p}]}^{sources\ and\ uptake\ by\ cells} \ in\ \Omega \tag{22}$$

Where:

- $\vec{p}$ is the vector of substrate densities.
- $\vec{p}^*$ is the vector of saturation densities.
- $\vec{D}$ is the vector of diffusion coefficients.
- $\vec{\lambda}$ is the vector of decay rates.
- $\vec{S}$ is the vector of Supply rates, which can vary across microenvironment.
- $\vec{U}$ is the vector of Uptake rates, which can vary across microenvironment.
- **Cell with index k, $1 \leq$ k $\leq$ Cells(t):**
  - $\vec{x}_k$ centre of the cell.
  - $\vec{W}_k$ volume of the cell.
  - $\vec{S}_k$ supply rate of the cell.
  - $\vec{U}_k$ uptake rate of the cell.
  - $\vec{p}^*_k$ saturation densities.

#### 1.4.2 Cell mechanics and movement

The mechanical interactions between cells are governed by the equations presented below. Cell movement can be categorized into two primary types: cell-cell potential forces and cell motility. The potentials models the net mechanical force on a cell as the result of interactions with neighboring cells, reflecting a balance of intercellular forces. In contrast, the latter captures active cellular movement, which may be either stochastic (random) or deterministic, depending on the desired behavioral model. Mechanics unit cases uses these heuristics.

##### Cell-cell potentials

$$\mathbf{v}_i = \sum_{j \in \mathcal{N}(i)} \left( \overbrace{-\sqrt{c^i_{\text{cca}} c^j_{\text{cca}}} \nabla\phi_{1,R_{i,A}+R_{j,A}}(\mathbf{x}_i - \mathbf{x}_j)}^{\text{cell–cell adhesion}} - \overbrace{\sqrt{c^i_{\text{ccr}} c^j_{\text{ccr}}} \nabla\psi_{1,R_i+R_j}(\mathbf{x}_i - \mathbf{x}_j)}^{\text{cell–cell repulsion}} \right) \times$$
$$- \overbrace{c^i_{\text{cba}} \nabla\phi_{1,R_{i,A}}(-\mathbf{d}(\mathbf{x}_i) \cdot \mathbf{n}(\mathbf{x}_i))}^{\text{cell–BM adhesion}} - \overbrace{c^i_{\text{cbr}} \nabla\psi_{1,R_i}(-\mathbf{d}(\mathbf{x}_i) \cdot \mathbf{n}(\mathbf{x}_i))}^{\text{cell–BM repulsion}} + \mathbf{v}_{i,\text{mot}} \tag{23}$$

Where:

- $\mathbf{v}_i$: Velocity of cell $i$.
- $\mathcal{N}(i)$: Set of neighboring cells of cell $i$.
- $\mathbf{x}_i$: Position vector of cell $i$.
- $R_i$: Radius of cell $i$.
- $R_{i,A}$: Adhesion radius of cell $i$ (maximum adhesion distance).
- $\phi_{1,R}$: Cell–cell adhesion potential function with interaction range $R$.
- $\psi_{1,R}$: Cell–cell repulsion potential function with interaction range $R$.
- $c^i_{\text{cca}}$: Cell–cell adhesion coefficient for cell $i$.
- $c^i_{\text{ccr}}$: Cell–cell repulsion coefficient for cell $i$.
- $c^i_{\text{cba}}$: Cell–BM adhesion coefficient for cell $i$.
- $c^i_{\text{cbr}}$: Cell–BM repulsion coefficient for cell $i$.
- $\mathbf{d}(\mathbf{x}_i)$: Distance vector from the cell to the basement membrane at position $\mathbf{x}_i$.
- $\mathbf{n}(\mathbf{x}_i)$: Normal vector to the basement membrane at position $\mathbf{x}_i$.
- $\mathbf{v}_{i,\text{mot}}$: Motility contribution to the velocity of cell $i$.

**Motility**

Each cell can set its own persistence time ($T_{\text{per}}$), migration speed ($s_{\text{mot}}$), migration bias direction ($\mathbf{d}_{\text{bias}}$, a unit vector), and a migration bias $b$ that ranges from 0 (Brownian motion) to 1 (deterministic movement along $\mathbf{d}_{\text{bias}}$). The motility velocity $\mathbf{v}_{\text{mot}}$ is added to the current velocity.

The probability of changing motility direction over time is:

$$\text{Prob}(\text{change } \mathbf{v}_{\text{mot}}) = \frac{\Delta t_{\text{mech}}}{T_{\text{per}}} \tag{24}$$

The standard migration bias formula is:

$$\mathbf{v}_{\text{mot}} = s_{\text{mot}} \cdot \frac{(1-b)\boldsymbol{\xi} + b\mathbf{d}_{\text{bias}}}{\|(1-b)\boldsymbol{\xi} + b\mathbf{d}_{\text{bias}}\|} \tag{25}$$

This equation interpolates from random (Brownian motion, $b = 0$) to deterministic migration ($b = 1$).

Where:

- $\mathbf{v}_{\text{mot}}$: Motility velocity vector.
- $s_{\text{mot}}$: Cell's migration speed.
- $\mathbf{d}_{\text{bias}}$: Migration bias direction (unit vector).
- $b$: Migration bias coefficient, from 0 (random) to 1 (deterministic).
- $\boldsymbol{\xi}$: Random unit vector representing Brownian motion.
- $\Delta t_{\text{mech}}$: Time step for mechanical updates.
- $T_{\text{per}}$: Persistence time between motility updates.

**Update position**

Cell position is updated using second-order Adam's Bashfort discretization:

$$\mathbf{x}_i(t + \Delta t_{\text{mech}}) = \mathbf{x}_i(t) + \frac{1}{2}\Delta t_{\text{mech}} \left(3\mathbf{v}_i(t) - \mathbf{v}_i(t - \Delta t_{\text{mech}})\right) \tag{26}$$

Where:

- $\Delta t_{\text{mech}}$: Time step used for mechanical updates.
- $\mathbf{x}_i(t)$: Position of cell $i$ at time $t$.
- $\mathbf{v}_i(t)$: Velocity of cell $i$ at time $t$.
- $\mathbf{v}_i(t - \Delta t_{\text{mech}})$: Velocity of cell $i$ at the previous time step.

#### 1.4.3 Phenotype: cell volume

The cell cycle in PhysiCell is implemented using a set of heuristic rules. It tracks the dynamic evolution of fluid and solid volumes within both the cytoplasm and the nucleus, with their sum constituting the total cell volume. To model transitions between different cell cycle phases, a volume transition factor is defined, capturing the volumetric changes associated with cellular progression. Volume transition factors ($f_{CN}$ and $f_F$) for proliferate is 0.5 and for apoptosis is 0.

**Total fluid volume**

$$\frac{dV_F}{dt} = r_F\left(V_F^*(t) - V_F\right) \tag{27}$$

**Nuclear solid**

$$\frac{dV_{NS}}{dt} = r_N\left(V_{NS}^*(t) - V_{NS}\right) \tag{28}$$

**Cytoplasmic solid**

$$\frac{dV_{CS}}{dt} = r_C\left(V_{CS}^*(t) - V_{CS}\right) \tag{29}$$

**Cell progresses (division, death)**

$$V_{CS}^*(t) = f_{CN}V_{NS}^*(t) \tag{30}$$

$$V_F^*(t) = f_F V(t) \tag{31}$$

**Where:**

- $V_F$: Current fluid volume
- $V_{NS}$: Current nuclear solid volume
- $V_{CS}$: Current cytoplasmic solid volume
- $V_F^*(t)$: Target fluid volume
- $V_{NS}^*(t)$: Target nuclear solid volume
- $V_{CS}^*(t)$: Target cytoplasmic solid volume
- $V(t)$: Total cell volume
- $r_F$, $r_N$, $r_C$: Rate constants for fluid, nuclear, and cytoplasmic volume change
- $f_F$: Fraction of total volume that is fluid
- $f_{CN}$: Ratio of cytoplasmic to nuclear volume

### 1.5 TiSim

Information can be found in these papers [Hoehme and Drasdo, 2010,Hoehme et al., 2010,Hoehme et al., 2023].

#### 1.5.1 Diffusion

Change of concentration is modeled using:

$$\delta c(r,t)/\delta t = D\sum_{i=1}^{d}\frac{(\delta^2 c(r,t))}{(\delta r_i^2)} - \hat{\gamma}\hat{\Theta}(c)n(r,t) \tag{32}$$

**Where:**

- **D**: diffusion coefficient
- **r**: consumption rate
- $\hat{\Theta}$: is equal to 1 if c $\geq$ 0 and it value is 0 othwewise.

#### 1.5.2 Mechanics - Cell movement

Tisim models cell movement by using a motion equation for each individual cell that accounts for the balance of various forces, including friction with the environment and active micro-motility. The model incorporates several factors: cell-cell friction and adhesion, cell-substrate interactions, and the presence of external forces. In addition, friction tensors describe the resistance to movement between cells and their surroundings and include different coefficients for perpendicular and parallel interactions. The model also includes a stochastic term to simulate random movement, emphasizing the role of micro-motility in driving cell migration.

**Equation for Cell Velocity**

$$\bar{\zeta}_{i_{\mathrm{ECM}}}^{CECM}\vec{v}_i(t) = \sum_{j\in \mathrm{NN}i} \bar{\zeta}_{ij}^{CC}\left(\vec{v}_j(t) - \vec{v}_i(t)\right) + \sum_{j\in \mathrm{NN}i} \vec{F}_{ij}^{CC} + \sum_i \vec{F}_{i_{\mathrm{ECM}}}^{f} + \vec{F}_i^{\mathrm{ext}} + \sum_i \vec{F}_i^{\mathrm{active},C}. \tag{33}$$

Where:

- $\bar{\zeta}_{i_{\mathrm{ECM}}}^{\mathrm{CECM}}$ is the drag coefficient between the cell and the ECM.
- $\bar{\zeta}_{ij}^{\mathrm{CC}}$ is the drag coefficient between neighboring cells $i$ and $j$.
- $\vec{F}_{ij}^{\mathrm{CC}}$ is the intercellular force between neighboring cells.
- $\vec{F}_{i_{\mathrm{ECM}}}^{f}$ is the frictional force with the ECM.
- $\vec{F}_i^{\mathrm{ext}}$ is the external force acting on cell $i$.
- $\vec{F}_i^{\mathrm{active},C}$ is the active force generated by cell $i$.
- NN$i$ indicates the set of nearest neighbors of cell $i$.

**Cell-Cell interaction**

Tisim includes two models to aproximate cell interaction forces ($\vec{F}^{\mathrm{CC}}$).

**Hertz-model**

$$F_{ij}^{\mathrm{DMT}} = \left|F_{ij}^{\mathrm{eHertz}}(d_{ij})\right|, \tag{34}$$

where $d_{ij}$ is the distance between the centers of two interacting spheres $i$ and $j$. The elastic Hertzian contact force is given by:

$$F_{ij}^{\mathrm{eHertz}} = \frac{4\tilde{E}_{ij}}{3}\sqrt{\tilde{R}}\delta^{3/2} - \pi\tilde{\gamma}\tilde{R} \tag{35}$$

The terms in this equation are defined as follows:

- $\delta = R_i + R_j - d_{ij}$ is the overlap distance between cells $i$ and $j$, where $R_i$ and $R_j$ are their respective radii.
- $\tilde{R}$ is the effective radius defined by $\tilde{R}^{-1} = R_i^{-1} + R_j^{-1}$.
- $\tilde{E}_{ij}$ is the effective Young's modulus (elastic response of two interacting cells) defined by:

  $$\tilde{E}_{ij}^{-1} = (1-\nu_i^2)E_i^{-1} + (1-\nu_j^2)E_j^{-1},$$

  where $E_i$, $E_j$ are the Young's moduli and $\nu_i$, $\nu_j$ are the Poisson's ratios of cells $i$ and $j$, respectively.
- $\tilde{\gamma} \approx \hat{\gamma} = \varsigma_m W_s$ is the specific adhesion energy, where:
  - $\varsigma_m \approx 10^5/\mathrm{m}^2$ is the density of adhesion molecules,
  - $W_s \approx 15 - 25k_bT$ is the binding energy of a single molecular bond.

The adhesion term $\pi\tilde{\gamma}\tilde{R}$ approximates the energy due to cell-cell contact, assuming no additional deformation beyond the Hertz contact area.

**JKR-model**

$$F_{ij}^{\mathrm{JKR}} = \left|F_{ij}^{\mathrm{JKR}}(d_{ij})\right| \tag{36}$$

**Where:**

- $\mathbf{d}_{ij}$ is the distance between two interacting cells calculated from two implicit equations [Attard, Parker, 1992]:

$$\delta = \frac{a^3}{\tilde{R}} - \sqrt{\frac{2\pi\hat{\gamma}a}{\tilde{E}_{ij}}} \tag{37}$$

$$a^3 = \frac{3\tilde{R}}{4\tilde{E}_{ij}} \left[ F_{ij}^{JKR} + 3\pi\hat{\gamma}\tilde{R} + \sqrt{6\pi\hat{\gamma}\tilde{R}F_{ij}^{JKR} + (3\pi\hat{\gamma}\tilde{R})^2} \right] \tag{38}$$

**Where**

- **a** is the contact radious.
- $\tilde{R}$ is the effective radious defined by: $\tilde{R}^{-1} = R_i^{-1} + R_j^{-1}$.
- The effective (composite) Young's modulus $\tilde{E}_{ij}$ is defined as:

$$\tilde{E}_{ij}^{-1} = (1 - \nu_i^2)E_i^{-1} + (1 - \nu_j^2)E_j^{-1}$$

- The specific adhesion energy $\hat{\gamma} \approx \varsigma_m W_s$**, being** $\varsigma_m \approx \frac{10^5}{m^2}$ the density of adhesion molecules and

#### 1.5.3 Cell grow and division

During G1, S, and G2-phase (interphase) it is assumed that a cell doubled its volume, and in M-phase deforms into a dumb-bell at constant volume until division. A certain cell does not re-enter the cell cycle if the pressure exerted on it overcomes a threshold value. The decision of whether a cell enters into the cell cycle was now made in two steps:

- Sampling of candidate cells for cell cycle entrance and determining the acceptance modeling a Poisson process.
- A cell is allowed to enter the cell cycle only if its mechanical pressure $p_i$ is below a threshold $p_{th}$. If $p_i \geq p_{th}$, the cell remains inactive.

During interphase, the cell targets to grow as twice as its initial volume $V_{DIV} = 2V_0$. This condition marks the entrance of the mitosis phase. The duration of the phase can be determined determined or stochastically, and the cells are adhered forming a dumbell shape. Presure is computed using:

$$p_i = \frac{\sum_j (F_{ij}^{CX} u_{ij})}{A_{ij}} \tag{39}$$

**Where**

- $F_{ij}^{CX}$ denotes the interaction force between cell i and object j (eg.floor of a preti dish).
- $u_{ij}$: normal vector pointing from cell i to object j.
- $A_{ij}$: interface between the cell i and object j.
